# Physiological trait networks enhance understanding of crop growth and water use in contrasting environments

**DOI:** 10.1101/2022.03.11.482897

**Authors:** Sean M. Gleason, Dave M. Barnard, Timothy R. Green, D. Scott Mackay, Diane R. Wang, Elizabeth A. Ainsworth, Jon Altenhofen, Timothy J. Brodribb, Hervé Cochard, Louise H. Comas, Mark Cooper, Danielle Creek, Kendall C. DeJonge, Sylvain Delzon, Felix B. Fritschi, Graeme Hammer, Cameron Hunter, Danica Lombardozzi, Carlos D. Messina, Troy Ocheltree, Bo Maxwell Stevens, Jared J. Stewart, Vincent Vadez, Joshua Wenz, Ian J. Wright, Kevin Yemoto, Huihui Zhang

**Affiliations:** Agricultural Research Service, United States Department of Agriculture, Water Management and Systems Research Unit, Fort Collins, CO 80526, USA; Department of Geography & Department of Environment and Sustainability, University at Buffalo, Buffalo, NY, 14261, USA; Department of Agronomy, Purdue University, West Lafayette, IN, 47907, USA; Agricultural Research Service, United States Department of Agriculture, Global Change and Photosynthesis Research Unit, Urbana, IL 61801, USA; Northern Colorado Water Conservancy District, Berthoud, CO 80513, USA; School of Natural Sciences, University of Tasmania, Hobart, TAS, 7001, Australia; Australian Research Council Centre of Excellence for Plant Success in Nature and Agriculture, The University of Tasmania Node, Hobart, TAS, 7001, Australia; Université Clermont Auvergne, INRAE, PIAF, 63000 Clermont-Ferrand, France; Queensland Alliance for Agriculture and Food Innovation, The University of Queensland, Brisbane, QLD 4072, Australia; Australian Research Council Centre of Excellence for Plant Success in Nature and Agriculture, The University of Queensland Node, St. Lucia, QLD, Australia; Université Bordeaux, INRAE, BIOGECO, Bât. B2, Allée Geoffroy St-Hilaire, CS50023, F-33615, Pessac cedex, France; Division of Plant Science and Technology, University of Missouri, Columbia, MO 65211, USA; Department of Biology, Colorado State University, Fort Collins, CO 80523, USA; National Center for Atmospheric Research (NCAR), Climate & Global Dynamics Lab, Boulder, CO 80305, USA; Horticultural Sciences Department, University of Florida, Gainesville, FL 32611, USA; Department of Forest and Rangeland Stewardship, Colorado State University, Fort Collins, CO 80523, USA; ICRISAT, Patancheru 502324, Andhra Pradesh, India; Hawkesbury Institute for the Environment, Western Sydney University, Locked Bag 1797, Penrith, NSW 2751, Australia; Department of Biological Sciences, Macquarie University, North Ryde, NSW 2109, Australia; Australian Research Council Centre of Excellence for Plant Success in Nature and Agriculture, Western Sydney University Node, Penrith, NSW 2751, Australia

**Keywords:** maize, plant growth, hydraulic traits, xylem, stomata, water potential, photosynthesis, crop improvement, breeding, process simulation

## Abstract

Plant function arises from a complex network of structural and physiological traits. Explicit representation of these traits, as well as their connections with other biophysical processes, is required to advance our understanding of plant-soil-climate interactions. We used the Terrestrial Regional Ecosystem Exchange Simulator (TREES) to evaluate physiological trait networks in maize. Net primary productivity (NPP) and grain yield were simulated across five contrasting climate scenarios. Simulations achieving high NPP and grain yield in high precipitation environments featured trait networks conferring high water use strategies: deep roots, high stomatal conductance at low water potential (“risky” stomatal regulation), high xylem hydraulic conductivity, and high maximal leaf area index. In contrast, high NPP and grain yield was achieved in dry environments with low late-season precipitation via water conserving trait networks: deep roots, high embolism resistance, and low stomatal conductance at low leaf water potential (“conservative” stomatal regulation). We suggest that our approach, which allows for the simultaneous evaluation of physiological traits and their interactions (i.e., networks), has potential to improve crop growth predictions in different environments. In contrast, evaluating single traits in isolation of other coordinated traits does not appear to be an effective strategy for predicting plant performance.

**Summary statement:** Our process-based model uncovered two beneficial but contrasting trait networks for maize which can be understood by their integrated effect on water use/conservation. Modification of multiple, physiologically aligned, traits were required to bring about meaningful improvements in NPP and yield.

## Introduction

Given the challenge to feed an increasing human population in the face of climate change, the need for improved crop genotypes has never been more important (Ainsworth and Ort 2010; Flörke et al. 2018; Hasegawa et al. 2018; Bailey-Serres et al. 2019; IPCC 2021). However, current efforts to improve crops are beset by immense systems complexity – nearinfinite combinations of soil, climate, plant, and management interactions (Spiertz et al. 2007). Although experimental methods in isolation have little chance to evaluate the scale of this complexity in a meaningful way, the integration of experimental methods and results with modeling represents a possible way forward for assessing trait combinations and their consequences on crop performance (Hammer et al. 2002).

Explicit representation of key biotic and abiotic processes is essential to develop a predictive understanding of plant function and the interactions between plant, climate, and soil (Holzworth et al. 2014; Mackay et al. 2015). Mechanistic plant models (i.e., process-based models) have therefore often been used to explore crop management strategies (Zhao et al. 2015), physiology by climate interactions (Bauerle et al. 2014), physiological trait coordination (de Wit 1965; Gifford et al. 1984), climate change impacts (Peng et al. 2020) and, more recently, trait selection, i.e., in combination with gene-to-phenotype trait models (Messina et al. 2009, 2018; Technow et al. 2015; Hammer et al. 2019; Wang et al. 2019; Cooper et al. 2021). Mechanistic models appear particularly well-suited to evaluate combinations of structural, morphological, and physiological traits, provided that key traits (and their interactions) are represented accurately (Alam et al. 2014; Sperry et al. 2016). However, there remains much uncertainty about which trait combinations are desirable in specific contexts and how much biological complexity is needed in models, given the breadth of applications (Hammer et al. 2019; Peng et al. 2020; Cooper et al. 2021).

Mechanistic models must simulate hypothetical trait networks of interest, i.e., the appropriate mechanisms and interactions relevant for the question being asked (Di Paola et al. 2016). Here, we focus on identifying the key interactions among physiological processes that control carbon-water exchange, and how these interactions manifest as differences in growth and yield in contrasting climates (Fig. 1). Given the complexity of the traits involved (e.g., photosynthesis, stomatal conductance, xylem water transport) and the heterogeneity of possible production environments (known as the target population of environments; TPE), we expected that key insights would be learned from the emergent behavior of the model itself, in addition to the outcomes of hypotheses testing. Key to this approach is our assumption that the explicit representation of water-carbon linkages would allow for a more predictive understanding of trait interactions and how traits could be manipulated in theory (e.g., via breeding programs) to improve crop growth and grain yield across the TPE.

**Fig. 1.**
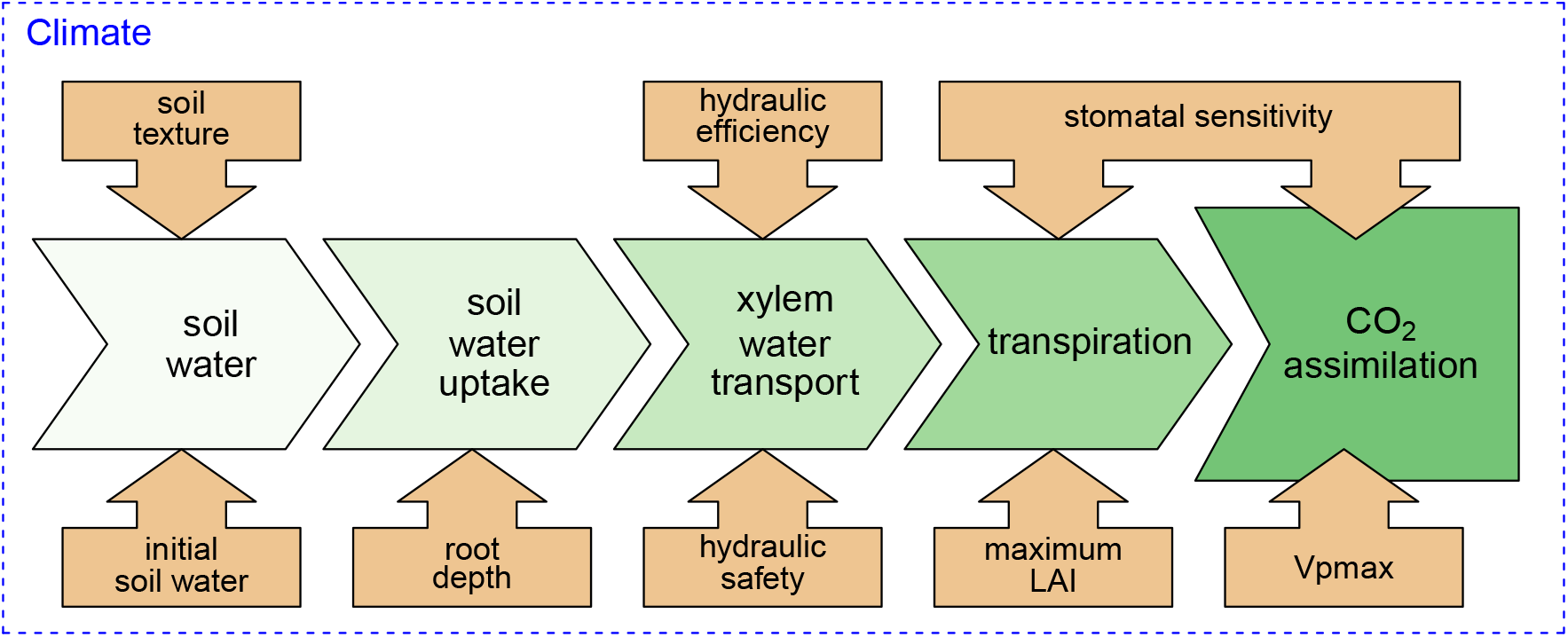
Key interactions among physiological processes (green shaded arrows) that control carbon-water exchange, and how these interactions manifest as differences in CO_2_ assimilation in a given climate (blue box). Traits manipulated in this study are represented by beige shaded arrows.

The exchange of water for atmospheric CO_2_ depends critically on the plant vasculature to deliver water to the sites of evaporation in the leaves (Brodribb et al. 2007). However, large quantities of water (200 – 1100 g) are required to obtain a single gram of CO_2_ (Shantz and Piemeisel 1927). As such, the conductive capacity of the vasculature needs to be closely coordinated with stomatal conductance and photosynthesis (Brodribb et al. 2017; Martin-StPaul et al. 2017; Deans et al. 2020; Xiong and Nadal 2020). However, transporting water long distances within plants cannot be done without risk because water is drawn through narrow xylem conduits (vessels and tracheids) in a metastable state under negative pressure. As the water content of the soil and atmosphere decrease, the negative pressure inside these conduits also decreases. If the pressure becomes too low, tiny bubbles of gas are pulled into the xylem, where they rapidly expand and block the conduits. These “cavitated” or “embolized” conduits are thereafter nonfunctional unless they can be refilled or replaced. As more conduits become embolized, the potential photosynthetic yield of the plant drops (Gleason et al. 2017b; Cardoso et al. 2018), or in severe cases, leaf tissue becomes damaged (Brodribb et al. 2021) and the risk of whole plant hydraulic failure increases (Meinzer and McCulloh 2013).

Given that gas exchange and growth depend critically on water transported via the xylem *and* this process is vulnerable to failure, many physiological-based plant growth models include hydraulic representation (Mackay et al. 2015; Venturas et al. 2018; Kennedy et al. 2019; Mencuccini et al. 2019; Danabasoglu et al. 2020; Cochard et al. 2021). Here, we used a modified version of one such model, the Terrestrial Regional Ecosystem Exchange Simulator (TREES) (Mackay et al. 2015, 2020) to evaluate structural and physiological processes, and how they interact in trait networks to govern the uptake, transport (soil-to-leaf), and exchange of water for CO_2_ in maize (*Zea mays*) grown under contrasting soil and climate conditions (Fig. 1). We addressed two questions: 1) can manipulation of the soil-to-atmosphere continuum via root, xylem, and stomatal traits confer improved growth and yield under water limitation? 2) what are the key plant traits and their bio-physical interactions that result in improved growth and yield under diverse climate scenarios?

## Methods

### Coupled hydraulic–carbon model (TREES)

TREES has been used to successfully simulate hydraulic-carbon dynamics in gymnosperms (Mackay et al. 2015, 2020) and angiosperms (Wang et al. 2020a), including maize (Mackay et al., in review). The published references cited above provide a more detailed description of the model, as well as examples of TREES model validation. Here we describe the basic features of the model, its parameterization (for maize), and further validation for field grown maize using a sap flow dataset. Hydraulic-carbon coupling is represented in TREES by integrating soil-xylem conductivity (Sperry et al. 1998; Mackay et al. 2015), Penman-Monteith energy balance (Monteith and Unsworth 1990), C_4_ photosynthesis (von Caemmerer 2013), and carbon allocation (Mackay et al. 2015, 2020) sub models. Key parameter settings (“static” parameters) and manipulated traits (“dynamic” parameters) are given in Table 1.

**Table 1.**
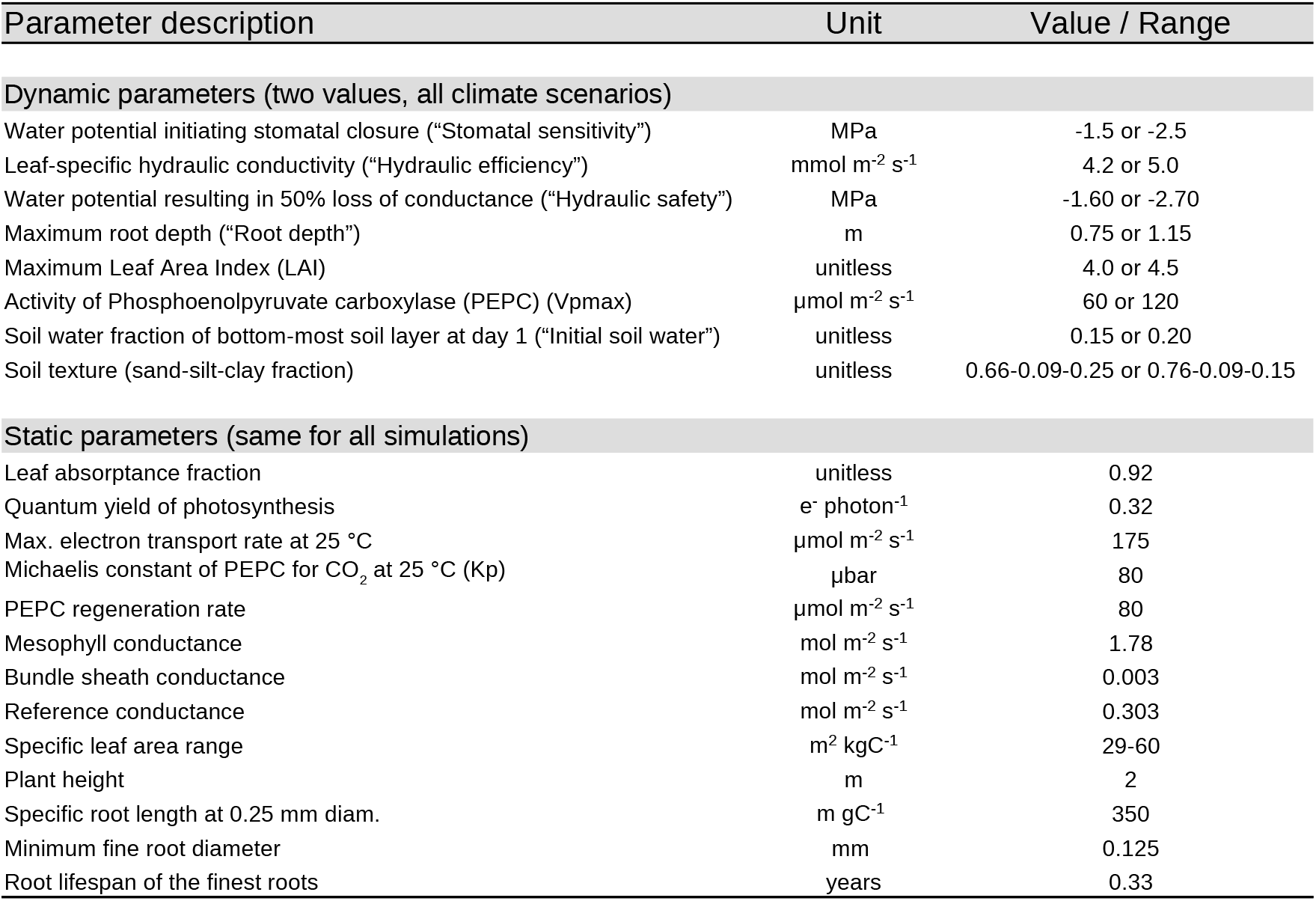
Parameter settings for all manipulated traits (“dynamic parameters”) and for parameters that were constant across all simulations (“static” parameters). Specific leaf area (SLA) was calculated as a function of the net CO_2_ assimilation rate and the amount of stored carbon (starch), and was allowed to vary within the given range.

Soil water uptake into roots is calculated as a function of root area, soil and root conductivity, and the driving force (water potential difference between root and soil) for each of five horizontal soil layers. The number of soil-root layers can be specified by the user. Soil conductivity (between root and bulk soil) and the conductivity of each root, stem, and leaf xylem segment is obtained via integral transformation of the Richards’ equation. Richards’ equation is a nonlinear partial differential equation that represents the unsaturated movement of water in soils and which, except in simple cases (e.g., uniform soil), has no analytical solution. TREES divides the root-soil interface into discrete “shells” of increasing distance from the root, and estimates flow within each shell using the Kirchoff transform, which allows for accurate water flow estimates (< 2% error in most cases) in heterogeneous soil using a mass-conservative “mixed-form” of the Richards’ equation (Ross and Bristow 1990; Sperry et al. 1998). Bulk water movement between soil layers is calculated via iteration of Darcy’s law.

Initial maximum whole-plant hydraulic conductance per unit leaf area was based on midday sap flow measurements taken on mature maize plants and predawn and midday leaf water potentials (Han et al. 2018). Embolism vulnerability was parameterized for each xylem segment (roots, stems, leaves) using vulnerability curves (2-parameter Weibull functions) obtained on field-grown maize stems (Gleason et al. 2019). Further information on how the embolism vulnerability curves were fit and interpreted is given in the supplemental materials (Fig. S1). At each 30-min modeled time-step, water movement, water potential, and xylem conductivity were determined via iterative solution, until a stability threshold was met or exceeded. Loss of xylem conductivity resulting from cavitation and embolism spread was remembered, allowing for progressive conductivity loss as xylem water potential declined. Although TREES allows for different Weibull coefficients for each root, stem, and leaf xylem segment, we used the same coefficients for all xylem segments, i.e., native embolism vulnerability was not allowed to differ among organs. It is likely that maize can generate positive pressure (ca. 0.14 MPa) in its root and stem xylem at night when soil water potentials exceed ca. −0.4 MPa (Gleason et al. 2017a). Considering it is unlikely that embolism in roots and stems could exist under positive pressure, we allowed xylem conductivity to fully recover when soil water potential was greater than or equal to −0.4 MPa.

Stomatal conductance was first estimated following the Whitehead-Jarvis application of Darcy’s law (including soil and xylem conductivity) to plant canopies (Whitehead 1998). TREES was modified in this study to allow for “conservative” and “risky” water use strategies by reducing stomatal conductance as a function of leaf water potential via an inverse logit model. This allowed for manipulation of the hydraulic “safety factor” (relationship between stomatal conductance and xylem water potential) via the midpoint and rate coefficients. Coefficient values were based on measurements made on field-grown maize plants (Gleason et al. 2021). Stomatal conductance was not allowed to decline below a minimum “cuticle” conductance (g_min_) value, set to 3.05 mmol m^-2^ s^-1^ based on greenhouse grown maize plants (Gleason et al. 2017b). Transpiration was then calculated from stomatal conductance via Penman Monteith energy balance and used to update the soil-xylem hydraulics (Mackay et al. 2015).

Net CO_2_ assimilation (A_net_) was calculated using the von Caemmerer C_4_ photosynthesis model (von Caemmerer 2013), which considers both enzyme limitation (e.g., when internal CO_2_ [C_i_] is saturating), as well as electron transport limitation (e.g., when irradiance is low). Temperature-dependent enzyme activities were modeled with Arrhenius functions (von Caemmerer 2013). The photosynthesis model was parameterized using A_net_~C_i_ measurements made on mature field-grown maize (Leegood and von Caemmerer 1989; Markelz et al. 2011; Gleason et al. 2017b).

Carbon allocation to roots, stems, and leaves was controlled by both carbon supply (photosynthesis) and hydraulic limitation (embolism). Leaf area index (LAI) was increased/decreased as the carbon available for growth and the specific leaf area (SLA; fresh leaf area divided by leaf carbon mass) increased/decreased. SLA was re-calculated at each time step and was calculated as a function of net CO_2_ assimilation rate (Wright et al. 2004) and the amount of stored carbon (starch). Root carbon was allocated to each of the five soil-root layers partially depending upon the hydraulic status of each layer, with larger carbon fractions allocated to more hydrated layers. After the vegetative growth stages were complete, the model shifted the allocation of non-structural carbohydrates to reproductive structures, e.g., grain development. All computations were done on 30-minute time steps.

### Validation of TREES for maize

TREES has been previously validated for maize using field datasets collected in 2012 and 2013 at the USDA-ARS Limited Irrigation Research Farm in Greeley, Colorado (40.4486 latitude, −104.6367 longitude, 1426 m elevation) (Mackay et al., in review). This includes validation against field measurements of leaf area index (LAI), sap-flow (whole-plant transpiration), soil water content by soil layer, and leaf water potential. Additionally, we provide further validation here using an additional sap flow dataset collected in 2017 from the same site (Greeley, Colorado). Sap-flow was measured using energy balance sensors (i.e., “heat pulse”) and sapIP dataloggers (Dynamax, Inc, Houston, TX, USA). Two sap flow sensors were placed on two representative plants selected randomly from within fully watered and water limited treatments. Fully watered treatments replaced 100% of unstressed crop evapotranspiration (ET) via irrigation and rainfall, whereas water limited treatments supplied 40% of unstressed crop ET. Plants were located within 20 m of one another and sap flow sensors were installed as described in Han et al. (2018). Data were collected from July 26 to September 7, 2017. sap flow simulations used 30-min mean values for precipitation, air temperature, wind speed, relative humidity, total shortwave radiation, and photosynthetically active radiation. Data were downloaded from a weather station (Station GLY04; Colorado Agricultural Meteorological Network) positioned within 50 m of the planted maize crop surrounded by trimmed and well-watered grass (reference conditions). Daily and seasonal variation in measured whole-plant transpiration (30-minute intervals) was well predicted by TREES. The fully watered treatment R^2^, residual standard error (RSE), and bias were 0.58, 0.655 kg m^-2^ d^-1^, and −0.381 kg m^-2^ d^-1^, whereas under limited water, the values of these fit statistics were 0.63, 0.522 kg m^-2^ d^-1^, and 0.111 kg m^-2^ d^-1^. Thus, TREES resulted in slightly more error and bias (negative bias; underestimated transpiration) under fully irrigated conditions than under limited water (Fig. S2).

### Simulation experiments

We evaluated the efficacy of physiological and structural trait combinations for two contrasting regions where maize is an important agronomic crop – the temperate (hot summer) climate of northeastern Missouri and the arid cold steppe climate of northeastern Colorado (Köppen-Geiger climate classification) (Beck et al. 2018). All simulations were run from June 1^st^ to November 9^th^. Parameter settings (traits) that were manipulated for the simulations are discussed individually below and key parameter settings are given in Table 1.

Twenty-years of meteorological data (ca. 2000 – 2020) were obtained from the University of Missouri, Missouri Historical Agricultural Weather Database (Knox County, MO) and the Colorado Agricultural Meteorological Network (Yuma County, CO). A typical “wet” year was chosen from the Missouri database as the year most closely aligned with the 75^th^ mean annual precipitation percentile. The total precipitation over the growth season (June 1 – November 9) for this scenario was 743 mm and included large early season precipitation events, followed by a relatively dry summer and large precipitation events occurring after September 25 (Hu and Buyanovsky 2003) (Fig. 2, “Central Plains Wet”). Considering that the amount and timing of precipitation is known to interact with other climate features (e.g., vapor pressure deficit; VPD) (Yuan et al. 2019), we focused on the effects of precipitation on plant growth by artificially creating a “dry” year for this site whilst conserving all other meteorological variables. This was done by reducing every precipitation event by 40%, giving a total seasonal precipitation for this scenario of 446 mm. Similarly, a typical “dry” year was chosen from the Colorado database as the year most closely aligned with the 25^th^ mean annual precipitation percentile. Total seasonal precipitation for this scenario was 289 mm, with most of this precipitation being received within the first 90 days of growth (Fig. 2, “High Plains Dry”). We then increased every precipitation event by 100% to create a wet season for this site (578 mm precipitation), keeping all other climate variables the same. In addition to simulating “wet” and “dry” years for both locations, we included a fully irrigated scenario for the Central Plains. For this scenario, we set the precipitation to zero and added 36-mm irrigation events every 3 days (hereafter, “irrigated”) (Fig. 2, “Irrigated”). We note that our manipulated climates (Central Plains Dry, High Plains Wet, Irrigated) are not meant to represent current or future climates at these locations but have been designed with the aim of achieving a better understanding of how traits might interact with precipitation at sites with contrasting VPD and temperature. Also, the labels “wet” and “dry” should not be viewed as a precipitation dichotomy because each climate scenario represents a different precipitation regime (amount and timing).

**Fig. 2.**
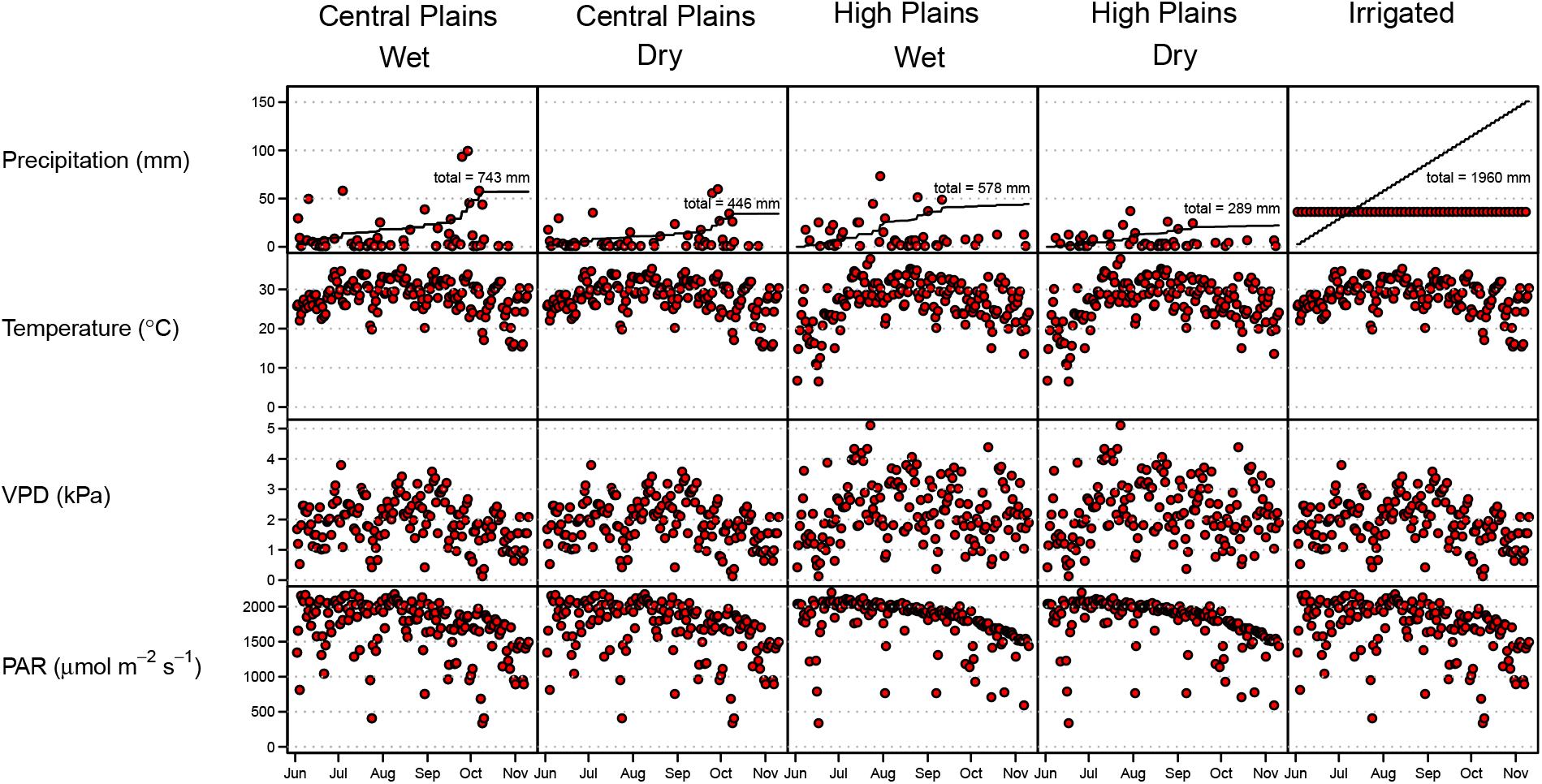
Daily precipitation, air temperature, vapor pressure deficit, and photosynthetically active radiation for each of the five climate scenarios. Cumulative precipitation for each climate scenario is represented with an unbroken black line.

Soil water holding capacity was manipulated by altering the soil textural properties of the whole soil column (Rawls and Brakensiek 1985; 1992). The sand-silt-clay fractions for the “fine” soil were set to 0.66-0.09-0.25, respectively, whereas these fractions for the “coarse” soil were set to 0.76-0.09-0.15. These modifications of soil texture resulted in water holding capacities of 25% for the fine soil and 18% for the coarse soil. In addition to manipulating the soil water holding capacity, we manipulated the starting value of the soil water content (volumetric fraction) for the bottom-most soil layer (0.75 m – 1.15 m), such that this layer was either “full” to field capacity (0.20 water fraction) or “not full” (0.15 water fraction) at the start of the growth season. The intention of this manipulation was to evaluate the shift in beneficial trait networks when deep antecedent soil water was readily available versus when it was limited.

Two levels of xylem embolism resistance were considered based on previous vulnerability curves constructed for maize (Gleason et al. 2017b, 2019, 2021). Whole-plant embolism resistance (all xylem segments) was simulated by setting the rate (b) and midpoint (c) Weibull coefficients. For embolism susceptible xylem, the rate and midpoint coefficients were set to 1.9 and 2.1, respectively (P_50_ = −1.6 MPa; P_88_ = −2.7 MPa), whereas the rate and midpoint coefficients were set to 2.7 and 2.1 (P_50_ = −2.3 MPa; P_88_ = −3.8 MPa) for embolism resistant xylem (Fig. S1). Whole-plant leaf-specific hydraulic conductance (hereafter “hydraulic efficiency”) was manipulated by setting it to either 0.104 or 0.124 g m^-2^ s^-1^ MPa^-1^ (Tsuda and Tyree 2000; Gleason et al. 2017b; Han et al. 2018). The intention of this manipulation was to evaluate the effect of water transport capacity on trait network coordination in the different climate scenarios.

Two levels of stomatal response to leaf water potential were considered based on previously measured stomatal conductance and leaf water potential measurements (Gleason et al. 2021). Stomatal closure was initiated when leaf water potential fell below −1.5 MPa (“conservative”) or −2.5 MPa (“risky”) and the leaf water potential resulting in a 50% loss of stomatal conductance was set to either −2.0 MPa (“conservative”) or −3.5 MPa (“risky”). The intention of this manipulation was to evaluate the effect of stomatal regulation on water use, carbon assimilation, and crop performance.

Deep and shallow root systems were simulated by either allowing or prohibiting root growth into the deepest soil layer (0.75-1.15 m). Wide and narrow leaf area to root area ratios were simulated by setting the maximum leaf area index (leaf area per unit ground area) to either 4.0 or 4.5 (Comas et al. 2019). Photosynthetic functioning was manipulated by setting the maximal activity of phosphoenolpyruvate carboxylase (Vpmax) to either 60 or 120 μmol m^-2^ s^-1^, based on the range reported in previous studies on maize (Leegood and von Caemmerer 1989; Pfeffer and Peisker 1998; Markelz et al. 2011; Perdomo et al. 2016; Gleason et al. 2017b).

All treatment combinations (2 levels of each trait) for soil texture, initial deep soil water fraction, xylem efficiency, embolism resistance, root depth, stomatal sensitivity, Vpmax, and leaf area index were simulated within each of the five climate scenarios (“Central Plains Wet”, “Central Plains Dry”, “High Plains Wet”, “High Plains Dry”, “Irrigated”), giving a total of 1,280 simulations. All simulations were compiled using the GNU Compiler Collection (GCC) on Ubuntu Linux operating systems.

### Data analyses

Treatments and treatment combinations were evaluated for each climate scenario using three approaches. Firstly, the efficacy of single traits was evaluated by determining the differences in mean seasonal net primary productivity (NPP; gross primary productivity minus respiration) and grain yield when the trait contrast was “high” versus “low” (e.g., high or low hydraulic efficiency), relative to the efficacy of all other traits. This was done by generating an ensemble of 350 decision trees using the randomForest package for R (Liaw and Wiener 2002). Each tree was created by sampling with replacement from the training dataset (50% of the dataset). Branch points at each node were resolved using a random subset of predictors. Over-fitting the training data was avoided in this way because each tree was fit with a different subset of simulations. “Importance” values for the decision trees were calculated for every trait as the reduction in model variance (unaccounted for variance in NPP or yield) when traits were included versus when they were omitted from the model. Thus, a high importance value means that including a particular trait in the decision tree model (e.g., manipulation of hydraulic safety to either a high or low value; Table 1) resulted in a meaningful increase/decrease in NPP or yield that was predicted by the model. Median, 25^th^ percentile, 75^th^ percentile, minimum, and maximum importance values were then calculated and used to evaluate single trait effects. To evaluate the interaction between time and individual traits, we plotted NPP and grain yield against the annual day (days since January 1^st^). This was done to determine if particular traits were more effective during specific periods of the growing season (e.g., early versus late season performance).

Considering that plants operate as connected trait networks, we focused our analyses on multiple trait effects, rather than single trait effects (Table 1 “dynamic parameters”). Therefore, our second analysis evaluated 2-trait effects by plotting all 2-trait combinations as heatmaps using the pheatmap package in R (Kolde 2019). Pheatmap is a hierarchical clustering and mapping function that allowed us to visually represent the mean effect of every possible two trait combination (e.g., conservative stomata + deep roots) on NPP and grain yield within each climate scenario. This provided a quick and intuitive representation of the best and worst performing two-trait combinations. Lastly, we expanded our random forest modeling to include up to four trait combinations. Decision tree models were fit to training datasets, created as described above, and then used to predict either NPP or grain yield in the test dataset. Specifically, 350 decision trees were fit for each climate scenario with each tree trimmed to four nodes (e.g., Vpmax → root depth → max LAI → gs sensitivity). An aggregate decision tree was then constructed for each climate scenario using the ctree and ggCtree (modified) packages for R (Hothorn et al. 2015; Martinez-Feria 2018). This method gives a robust analysis of the best trait combinations conferring improved performance in each climate scenario. When viewed in the context of individual trait effects and the timing of these traits throughout the growth season, these trait combinations provided information about why and when particular trait combinations were effective. These aggregate decision trees were also useful for evaluating multiple trait strategies in the contrasting climates. For example, they helped address the question: do we require specific trait combinations for each individual climate scenario, or are there some trait combinations that are likely to perform well across climates?

All data analyses and graphics were done using R software (R Core Team 2021). All data and code (R, C++) used in this study are in the public domain and can be downloaded from GitHub (https://github.com/sean-gl/trait_network_ms_TREES_data_and_code).

## Results

### Single trait effects

Efficacy of single traits differed markedly by climate scenario. High NPP and high yield simulations in both wet climate scenarios, with higher annual precipitation and sufficient late season precipitation, featured traits contributing to enhanced soil water extraction, efficient water transport, and high rates of gas exchange (deep roots, high hydraulic efficiency, high hydraulic safety, and risky stomata) (Figs 3, S3, S5). Similar traits were effective in conferring improved NPP and yield in the Central Plains Dry site, with the notable exception that conservative stomata (closure at higher water potential) were beneficial during late season growth (ca. after day 240), particularly during grain development (Fig. 4a). This result reflects the importance of achieving coordinated liquid- and gas-phase conductance when water is abundant, as well as traits conferring water conservation when water is scarce. Water conservation traits, access to deep soil water, and high instantaneous water use efficiency (conservative stomata, deep roots, high Vpmax) improved plant performance in the High Plains Dry scenario by reducing the adverse impact of late season water deficit (Figs 3, 4b, S6). In contrast to the three non-irrigated scenarios, irrigation kept soil water potentials near zero throughout the growing season, resulting in sufficient xylem water transport to support high rates of photosynthesis (even when hydraulic efficiency was low) with little risk of embolism, and thus featured traits maximizing canopy-level carbon income (high maximal LAI, high Vpmax) (Figs 3, S7). These three contrasting trait networks reflect the importance of a coordinated trait response that balances the canopy water demand with, not only soil water availability, but also the capacity to move this water through the xylem.

**Fig. 3.**
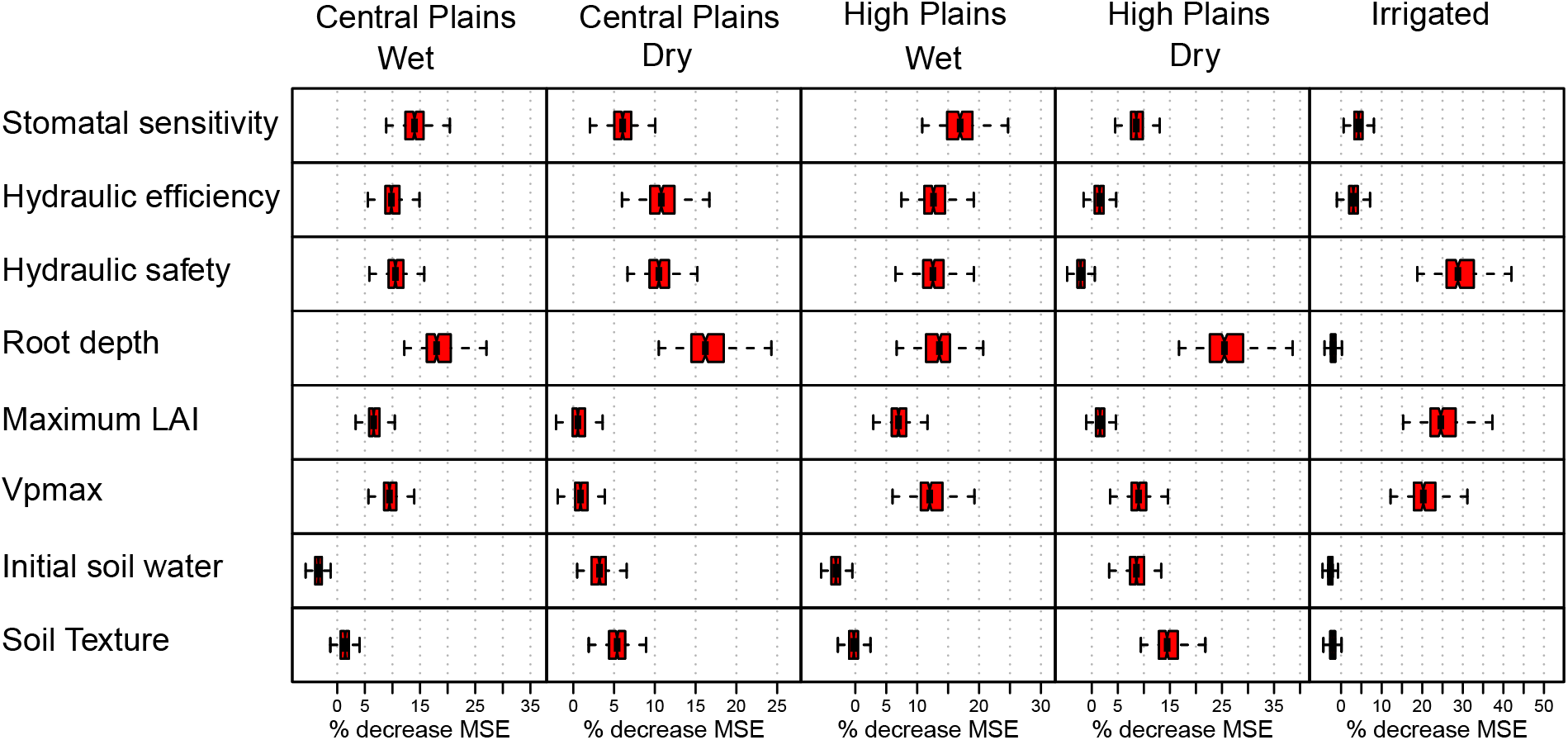
“Importance” scores for individual traits that have been derived from 350 decision tree ensembles. Larger importance values denote trait contrasts (e.g., deep vs shallow roots) that resulted in large differences in net primary productivity (NPP), i.e., reduction in root mean square error (MSE; square root of model variance) when the trait was included in the model. Small importance values reflect trait contrasts that resulted in smaller reductions in model variance.

**Fig. 4.**
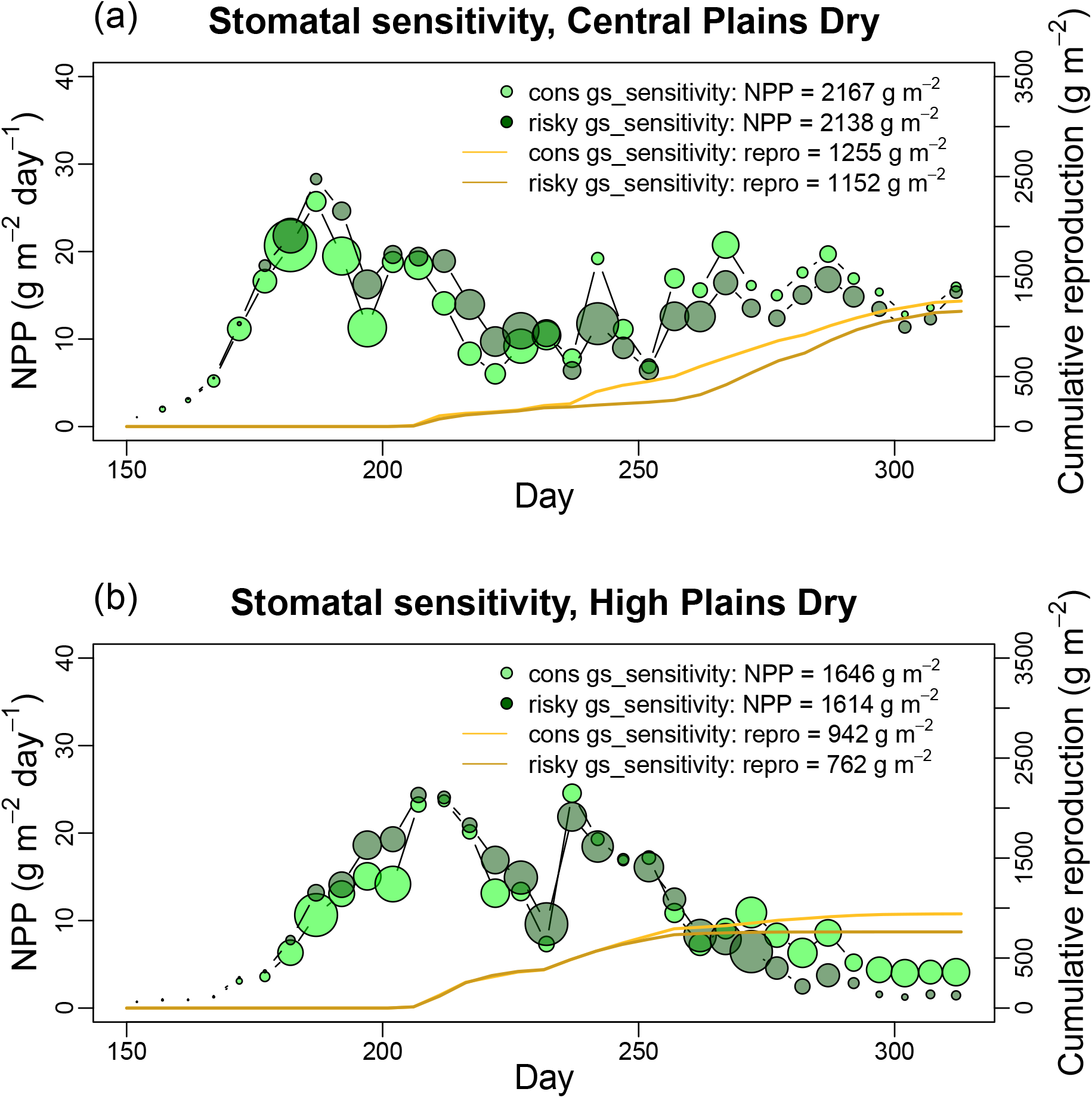
Net primary productivity (NPP) and reproductive output for stomatal sensitivity trait contrast (“risky” vs “conservative” stomatal response to leaf water potential) for the Central Plains, Dry (a) and the High Plains, Dry (b) sites. Symbol size has been scaled proportionately with variance in NPP across simulations.

Traits that were beneficial in the High Plains were generally also beneficial in the Central Plains, but there were notable exceptions to this pattern. Firstly, early season aboveground and belowground growth (ca. first 50 days of growth) was markedly faster in the Central Plains than in the High Plains, in both the wet and dry scenarios (Figs S3-S6 & S8-S11 “root depth”). This outcome arose mainly from differences in soil and air temperature between the two sites – with lower early season temperatures at the High Plains site (means ± SDs of 13.2 ± 6.2 °C and 3.5 ± 3.4 °C, respectively) than at the Central Plains site (means ± SDs of 21.6 ± 4.7 °C and 6.8 ± 4.2 °C, respectively) (Fig. 2). Secondly, risky stomatal regulation, in combination with higher VPD at the High Plains site (Fig. 2), resulted in faster and more complete extraction of soil water before it could be evaporated from shallow soil layers. This resulted in a larger fraction of the received precipitation passing through plant stomata (hereafter “transpiration fraction”; T-fraction) at the High Plains site than at the Central Plains site under both wet and dry scenarios (Figs S13-S16 “stomatal sensitivity”). For example, the transpiration fractions of plants with risky stomata were about 3% higher in the Central Plains Dry scenario and 6% higher in the High Plains Dry scenario (Figs S14 & S16, “T-fraction” in the “stomatal sensitivity” panel). Predictably, the tradeoff associated with risky stomata was lower precipitation use efficiency (NPP per unit total received precipitation; PrUE), which was 4% lower in both dry scenarios (Figs S14 & S16, “PrUE” in the “stomatal sensitivity” panel). This indicates that although plants with risky stomata achieved higher water use, they used this water less efficiently (lower instantaneous and seasonally integrated water use efficiency) than plants with more conservative stomata.

High PEP-carboxylase efficiency (increase in A per unit C_i_; Vpmax) was beneficial in every scenario, but especially in the wet and irrigated scenarios (Figs 3 & S3-S7 “Vpmax”). This suggests that if it could be selected for, it would likely improve the performance of crops in nearly every case. However, it is important to note that mitochondrial respiration does not scale to Vpmax, but rather to Vmax (rubisco activity) in our C_4_ photosynthesis model (von Caemmerer 2013). Although we should expect general alignment between Vpmax and Vmax within and across species (Schlüter and Weber 2020), we only examined the independent effect on variation in Vpmax on plant performance and therefore cannot extrapolate to its likely impact on mitochondrial respiration or to instances when soil nitrogen limits enzyme synthesis. As such, high carboxylation efficiency should not be viewed as a “free lunch”, as also supported by the wide range in Vpmax within and across species (Pilon-Smits et al. 1991), i.e., natural selection has not maximized Vpmax in all cases.

Coarse soil texture had a similarly positive effect on plant performance in both dry climates (Figs S4 & S6 “soil texture”). This effect was largely an outcome of manipulating the soil texture of the entire profile, rather than only the deeper layers. Fine soil texture (high field capacity) at the surface, combined with frequent but low volume precipitation events, resulted in much of the precipitation being held close to the surface and subject to evaporation. Additionally, low precipitation in the dry climate scenarios, coupled with low matric potential of fine textured soils, resulted in very little saturated (soil matric potential ≥ 0) and unsaturated (soil matric potential < 0) flow out the bottom of the rhizosphere and a meaningful fraction of soil water being held at water potentials to low for uptake (Figs S9 & S11 “soil texture”). These conditions resulted in lower transpiration fractions in the fine textured soil (Figs S14 & S16 “T-fraction” in “soil texture” panel). Predictably, when rainfall was increased, the effect of soil texture was reversed such that plants growing in finer textured soil (higher field capacity) had access to more water and achieved improved growth and reproductive output (Figs S3 & S5 “soil texture”). We note that the soil texture effect in the dry scenarios would be less conspicuous, and even likely reversed, in a natural soil where layer silicate clays have been translocated to deeper horizons (Buol et al. 2011).

Although examining single traits gives us some indication of which traits might be beneficial in certain climate scenarios, this approach cannot inform us about why particular traits appear to be beneficial in some cases and not others. For example, high variation in the importance values (reduction in residual variance when individual traits are included in the decision tree) (Fig. 3) indicates that some traits were only beneficial in simulations that included biologically aligned traits, and when these traits were omitted from the decision tree the simulation performed poorly. To obtain a better understanding of trait synergies (beneficial parameter interactions), as well as the biological reasons for them, we examined multiple trait effects simultaneously.

### Multiple trait effects

Trait combinations that increased access to deep water, efficient and safe water transport to the leaves, and high stomatal conductance were associated with improved growth and yield in both wet climate scenarios and the irrigated scenario (Figs S18, S20 & S22). Importantly, much of the variation in NPP and yield that was accounted for in the random forest models was dependent upon specific trait combinations. Modifications of individual traits, either in isolation or in combination with other poorly aligned traits, resulted in little improvement in NPP or yield. For example, deep rooting was most beneficial at the Central Plains Wet site, but only in combination with traits that facilitated the efficient and safe movement of this water to the leaves (high hydraulic efficiency, high hydraulic safety) and exchange of water for CO_2_ (risky stomata, high Vpmax, high maximum LAI) (Figs S18 & S23). Beneficial trait networks in the two dry climate scenarios differed from one another depending on the total amount of precipitation and the timing of precipitation. The High Plains Dry scenario, with lower seasonal precipitation and markedly low late season precipitation, featured networks that included conservative stomata (firstly) in coordination with access to deep soil water (deep roots), and uninterrupted xylem functioning during periods of low water potential (hydraulic safety) (Figs 6, S26). In contrast, the Central Plains Dry scenario, with higher total and late season precipitation, featured traits conferring access to deep soil water (deep roots) in coordination with safe and efficient water transport, and then conservative stomata (Figs S19, S24). Thus, differences in the timing and amount of precipitation resulted in notable differences in trait coordination, but also remarkable similarities, at least within the two “wet” and two “dry” scenarios.

The two-trait analysis of the High Plains Dry scenario revealed that nearly every simulation that did not include both deep roots and conservative stomata were largely failures, whereas the late season precipitation events and lower evaporation at the Central Plains site allowed for other alternative, albeit less successful, trait networks, e.g., high LAI coupled with high hydraulic safety and conservative stomata (Figs S24 & S26). Notably, the benefit of high LAI in this scenario was reversed when g_min_ (minimum stomatal and cuticle conductance to water vapor) was increased from 3 mmol m^-2^ s^-1^ to 10 mmol m^-2^ s^-1^, suggesting that stomatal “leakiness” may be an important trait to consider for future trait networks (Barnard and Bauerle 2013; Blackman et al. 2019), particularly if stomatal leakiness increases at higher temperatures (e.g., under climate change), which has been reported for some species (Slot et al. 2021).

### Seasonal dynamics

Differences in plant performance between the High Plains and Central Plains can be largely understood from the different seasonal trajectories of precipitation and temperature (air and soil). Firstly, the efficacy of the conservative water use strategies (e.g., conservative stomata, high hydraulic safety), depended critically on ample early season precipitation and low late season precipitation. In contrast, high water extraction and transport strategies (e.g., risky stomata, high hydraulic efficiency) were most beneficial, in the face of cold early season temperatures, when soil water was available water during grain development. This switch in the importance of water conserving versus water using strategies can be seen in the seasonal NPP plots under both dry climate scenarios (Fig. 4 a & b). In both of these scenarios, risky stomatal response (initiating stomatal closure at low xylem water potential; dark green symbols in Fig. 4) resulted in higher NPP during the first few weeks of growth when soil water was available, but later in season, when shallow soil water was largely depleted and the reproductive structures were developing, conservative stomata (initiating stomatal closure at high xylem water potential) conferred a strong advantage, especially in reproductive output (Fig. 4 a & b). Similarly, high hydraulic conductivity conferred an early season advantage at the High Plains Dry site, but later in the season resulted in poorer performance (Fig. S6 “hydraulic efficiency”).

Trait combinations associated with success in all climate scenarios reflected the relative costs and benefits of: 1) accessing shallow and deep soil water, minimizing losses to saturated/unsaturated flow and evaporation (deep roots), 2) transporting water efficiently through the xylem at low water potential (high hydraulic efficiency and safety), 3) the effective use of soil water after it reached the leaves, avoiding high VPD conditions (conservative stomata), and 4) achieving high instantaneous water use efficiency (high Vpmax). Even seemingly subtle differences in air and soil temperature, the timing of precipitation, the frequency and volume of precipitation events, and soil water storage capacity, resulted in meaningful differences in beneficial trait combinations (e.g., Central Plains Dry versus High Plains Dry; Figs 5, S19, S24, S26).

**Fig. 5.**
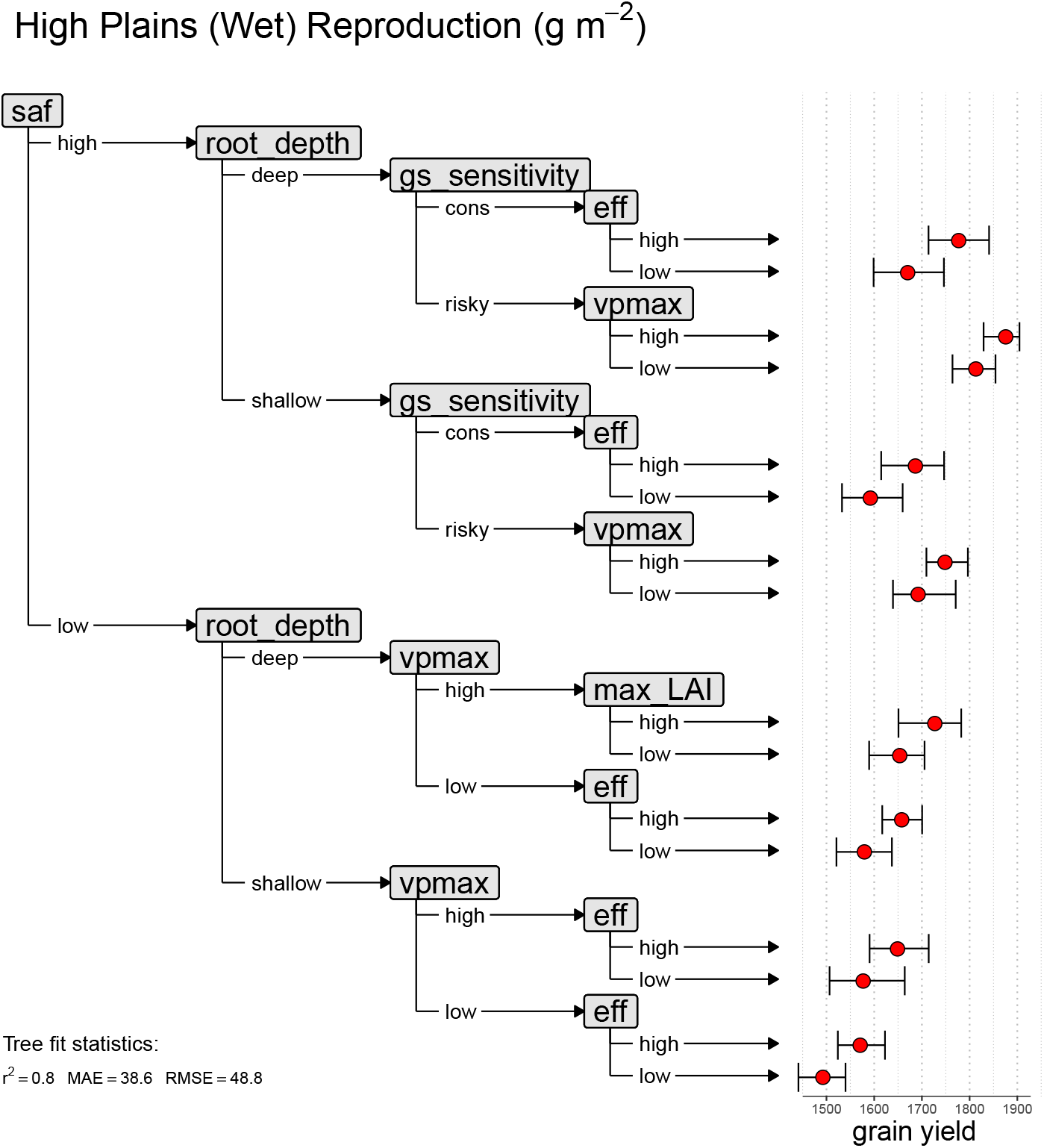
Representative multiple trait decision tree for the High Plains Wet scenario. Manipulated traits are represented with shaded boxes, whereas the contrasting values of these traits (e.g., “low”, “high”) are denoted by labeled arrows. The first branch point (trait contrast) is the trait resulting in the largest decrease in model variance, whereas the last branch point denotes the trait contrast resulting in the smallest decrease in model variance. The first four most important nodes (trait contrasts) are shown. Error bars denote +/- standard deviation (n=16). Vpmax = maximum activity of PEP-carboxylase. Root depth = maximum depth of root system. Max LAI = maximum achievable leaf area index. Saf =xylem embolism resistance. Gs sensitivity = stomatal response to leaf water potential. Eff = maximum xylem conductance.

**Fig. 6.**
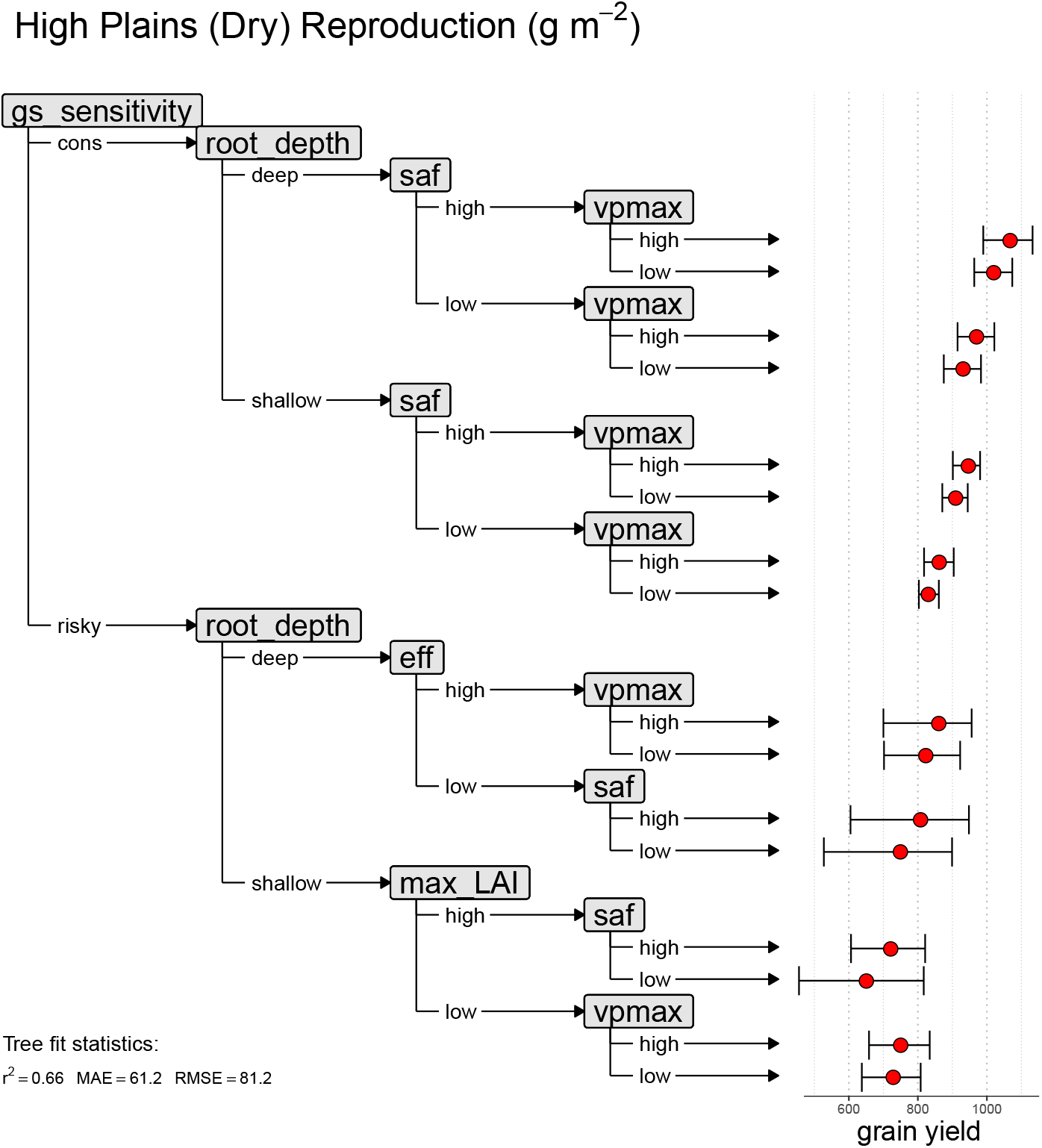
Representative multiple trait decision tree for the High Plains Dry scenario. Manipulated traits are represented with shaded boxes, whereas the contrasting values of these traits (e.g., “low”, “high”) are denoted by labeled arrows. The first branch point (trait contrast) is the trait resulting in the largest decrease in model variance, whereas the last branch point denotes the trait contrast resulting in the smallest decrease in model variance. The first four most important nodes (trait contrasts) are shown. Error bars denote +/- standard deviation (n=16). Vpmax = maximum activity of PEP-carboxylase. Root depth = maximum depth of root system. Max LAI = maximum achievable leaf area index. Saf = xylem embolism resistance. Gs sensitivity = stomatal response to leaf water potential. Eff = maximum xylem conductance.

## Discussion

The purpose of our simulations was to evaluate the potential efficacy of structural and physiological trait networks to improve the performance of maize grown under contrasting soil and climate conditions. It was not the purpose of our simulations to generate trait selection goals for any particular site or region of interest, and our results should be used with caution for this purpose. Thus, we place particular emphasis on biological interactions (trait combinations, rather than single traits) and the shift of these interactions across climates. However, simulating the outcomes of this complex biological system requires that we understand and can successfully model the important components of its complexity. In our case, we included eight soil, xylem, and leaf traits affecting soil water retention, soil water uptake, water transport to the leaves, and the exchange of water for atmospheric CO_2_ (Fig. 1). As such, our simulations represent an important network of traits governing the fluxes of water and carbon, and which exhibited coordinated shifts in their alignment to confer either high instantaneous CO_2_ uptake or soil water conservation, depending on the climate context. Although these simulated trait assemblages are hypothetical, they are supported by both empirical measurements and our conceptual understanding of plant functioning, in particular the well-understood linkages among water uptake (roots), water transport (xylem), stomatal conductance, and photosynthesis (Brodribb and Holbrook 2003; Brodribb et al. 2007; Creek et al. 2018; Deans et al. 2020), and the utilization of stored soil water during anthesis and ovule development (Sinclair et al. 2005; Vadez et al. 2014; Messina et al. 2015, 2021; Reyes et al. 2015; Diepenbrock et al. 2021).

### Effective seasonal transpiration trait networks

Traits leading to improved water availability during reproductive development in grain crops have been identified via comparative physiology and modeling studies, and include increasing transpiration efficiency (biomass produced per unit transpiration) by limiting maximal transpiration (Vadez et al. 2014; Messina et al. 2015), increasing net CO_2_ assimilation (Gilbert et al. 2011; Niinemets et al. 2017; Wang et al. 2020b), and reducing xylem conductivity (Richards and Passioura 1989; Sinclair et al. 2008; Choudhary and Sinclair 2014). Given that transpiration efficiency represents the integrated product of several structural and physiological traits (e.g., xylem-specific conductivity, xylem embolism resistance, stomatal regulation, root depth, and leaf/root surface area), the detailed modeling presented here allowed us to investigate the possible effects of these finer scale traits.

In our simulations we increased transpiration efficiency by either increasing the A~C_i_ slope (higher Vpmax) or else manipulating traits that resulted in reduced stomatal conductance, i.e., increasing the sensitivity of stomata to xylem water potential, reducing xylem conductivity, reducing xylem embolism resistance, or restricting root growth. Although higher PEP-carboxylase efficiency was associated with improved plant performance in all cases, the other trait manipulations resulted in reduced access to soil water (restricted root growth), or else slower relative growth rate (stomatal sensitivity, low xylem conductivity, low xylem safety) (Figs S3-S7 & S8-S12). Reducing xylem conductivity, either via lowering maximal conductivity or decreasing embolism resistance, did not result in meaningful improvements to growth in yield in the dry scenarios (Figs S19, S21, S24, S26). In contrast to this, increasing the stomatal sensitivity to leaf water potential resulted in markedly improved growth and yield (especially) in both dry climate scenarios (Figs S19, S21, S24, S26 “gs_sensitivity”). This difference in late season water conservation, resulting from lower xylem conductivity (not effective) versus higher stomatal sensitivity (effective), was unexpected because both these traits are functions of water potential. The reason for this difference was the timing of water use. In particular, “sensitive” stomata closed mainly during periods of low water potential (high VPD, midday hours), thus reducing midday transpiration, but also effectively preventing xylem embolism. The combined effect of this was improved deep soil water availability during grain development, higher precipitation use efficiency (Figs S14 & S16 “stomatal sensitivity”), improved water transport (without embolism), and effective gas exchange immediately after precipitation events (Figs S4 & S6 “stomatal sensitivity”, i.e., spikes in NPP after day 225). In contrast, reducing maximal hydraulic efficiency resulted in lower water use overall, but midday (high VPD) stomatal conductance and transpiration were higher than for the sensitive stomata trait. This resulted in lower daily and seasonally integrated water use efficiency (Fig. S14 & S16 “PrUE”).

The importance of fast early season growth, and especially early season root growth, is well aligned with previous empirical and simulated results (Tron et al. 2015; Palta and Turner 2019; Diepenbrock et al. 2021; Freschet et al. 2021). A recent analysis of 2,367 maize hybrids grown across 23 environments (North America and Chile) and 3 years found root elongation rate an important determinant of grain yield in combination with other structural and morphological traits (trait networks) (Diepenbrock et al. 2021). Similarly, the result reported here that high hydraulic efficiency was associated with improved performance in both dry climate scenarios also has empirical support (Gleason et al. 2019, 2021). For example, two maize field experiments performed in Colorado under water deficit (Gleason et al. 2019, 2021) reported that maize plants with high hydraulic efficiency transpired a greater fraction of soil water than low efficiency plants, but were also able to “self regulate” (decrease hydraulic conductance) as water potential declined (Pammenter and Vander Willigen 1998) (Fig. S1). The loss of xylem conductivity at low water potential was made even more beneficial in our simulations because we allowed roots and stems to regain conductive capacity overnight if sufficient soil water was available (see methods) (Gleason et al. 2017a). Thus, maize plants with intrinsically high hydraulic conductance were also able to achieve a relatively high precipitation use efficiency (Figs S14 & S16 “hydraulic efficiency”). However, given that that embolism reversal has never been directly observed (e.g., using microCT or Optical methods) in maize *leaves* (or the leaves of any other species), and claims of embolism reversal in other species have been questioned (Cochard and Delzon 2013; Johnson et al. 2018), our assumption that xylem conductivity can be perfectly restored overnight could be overly optimistic. Another possibility is that reduced soil-plant hydraulic conductance may result from a reversible decrease in rhizosphere conductivity (Figs S13-S17 gray bars) (Bourbia et al. 2021).

Although the direct effects of reduced stomatal conductance during midday (i.e., when VPD is high) has been reported elsewhere (Zaman-Allah et al. 2011; Turner et al. 2014; Vadez et al. 2014; Condon 2020; Collins et al. 2021), the interactions evident in our results between stomatal regulation, rooting depth, temperature (beyond its effect on VPD), and embolism resistance have not been previously noted. However, the importance of trait networks is being increasingly recognized in both plant physiology and genetics (Gleason et al. 2018, 2019; Hammer et al. 2019, 2021; Momen et al. 2019; Peng et al. 2020; Cooper et al. 2021; Diepenbrock et al. 2021). By utilizing biologically realistic statistical models (e.g., structural equation modeling, Gleason et al. 2019; Momen et al. 2019; He et al. 2020), as well as process-oriented plant growth models (Mackay et al. 2015; Holzworth et al. 2018; Venturas et al. 2018; Cochard et al. 2021), it is now possible to evaluate the physiological and structural determinants of transpiration efficiency, as well as the interactions and tradeoffs associated with these traits.

### Water uptake, xylem transport, and photosynthesis trait networks

Trait networks conferring improved crop performance under the wet and irrigated climate scenarios included relatively well-understood theoretical (Deans et al. 2020) and empirically observed linkages between soil water access, water transport to the sites of evaporation in the leaves, and the exchange of water for atmospheric CO_2_ (Brodribb and Holbrook 2003; Brodribb et al. 2007; Brodribb and Jordan 2008; Vadez 2014; Scoffoni et al. 2016; Martin-StPaul et al. 2017; Xiong and Nadal 2020). Similar trait assemblages have been found in maize, sorghum, sugarbeet, sunflower, wheat, olive, and chickpea (de Wit 1958; Steduto et al. 2007; Zhu and Cao 2009; Hanks 2015; Zhao et al. 2018; Gleason et al. 2019, 2021; Klimešová et al. 2020; Pires et al. 2020). Although there were important differences between the High Plains Wet and Central Plains Wet scenarios, as noted above, deep rooting, risky stomata, safe and efficient water transport, high Vpmax, and high maximal LAI, were advantageous, but only when aligned as a network with one another (e.g., Fig. 5). This trait network reflects the biological linkage between water uptake → water transport → stomatal conductance, and → high carboxylation efficiency (A~C_i_ slope) (Fig. 1). In addition to this result arising from carbon-water linkage, there were two less intuitive results. Firstly, the trait network that conferred improved performance in the wet and irrigated scenarios was meaningfully different from the trait network that conferred improved performance in the dry scenarios. *As such, superior genotypes tailored for the wet scenario would be ill-designed for dry scenarios (and vice versa), and especially dry scenarios where late season growth requires stored soil water*. However, roots, stomata, and photochemistry are known to be significantly plastic in field grown maize (Gleason et al. 2017b; Schneider et al. 2020; Ding et al. 2021). Although we do not address trait plasticity here, we should almost certainly expect attenuation of adverse intrinsic trait effects via a coordinated plastic response. The second important finding was that, even under fully watered conditions, transporting water from the soil to the leaves is a risky biological process. This is evident from the efficacy of high embolism resistance in every scenario, as well as the negative impact of xylem embolism on maize growth and reproductive development (e.g., “tassel blasting”) (Gleason et al. 2017b, 2019; Dong et al. 2020). This result is supported by multiple measurements of maize embolism resistance, which by all accounts is low, i.e., half the xylem conductive capacity is lost at relatively high/hydrated water potential (ca. −2.6 to −1.4 MPa) (Cochard 2002; Li et al. 2009; Gleason et al. 2017a, b, 2019). Higher embolism resistance in the wet and irrigated scenario was especially beneficial when combined with traits maximizing the delivery of liquid phase water to the stomata – deep roots, risky stomata, and high hydraulic efficiency, i.e., embolism safety was not particularly beneficial in isolation.

### Implications for crop improvement

Selection of a plant growth model should be guided by the needs of the user (McMaster and Ascough 2011; Di Paola et al. 2016). In the case we present here, modeling physiological processes and their interactions resulted in growth and water use outcomes that were broadly aligned with field measurements; however, it remains an important question how much biological resolution can be added (e.g., organ-level, protein-level, gene expression) without losing upper-level functioning and rigor (Hammer et al. 2019; Peng et al. 2020; Tardieu et al. 2020). Although we do not address this topic at length here, we caution that modeling fine scale physiological processes should not be viewed as a necessary step towards crop improvement, or even towards achieving better biological understanding. Given the difficulty of developing “bottom-up” models that perform well at higher levels of biological organization, hybrid approaches that allow for the nesting of specific lower order processes within whole-plant ecophysiological models may represent an effective bridge between fine scale and coarse scale modeling approaches (Tardieu et al. 2020).

The application of detailed process-based physiological models to assist breeding efforts has recently been discussed at length elsewhere (Messina et al. 2018; Hammer et al. 2019; Wang et al. 2019; Cooper et al. 2021), but the key advantage provided by such models is to breakdown higher order processes (e.g., transpiration) into their constituent components (e.g., xylem conductivity, xylem embolism resistance, stomatal conductance, xylem pressure gradient), and connect these component traits to causal genetic variation. For example, the development of AQUAmax® (Pioneer Hi Bred International, Inc., Johnston) maize hybrids, which were initially targeted for the western corn belt of North America, represent a coupling of water conservation, photosynthesis, and carbon partitioning traits, and thus required the careful consideration of multiple physiological processes (Cooper et al. 2014a, b; Messina et al. 2020). Assuming that modeling processes at these finer scales can reliably simulate plant performance, and also assuming that component traits can be linked with their corresponding functional nucleotide polymorphisms, it is then possible to predict trait values from the genotype and select target genotypes with desired traits (Hammer et al. 2019; Messina et al. 2020; Cooper et al. 2021).

Despite the potential usefulness of physiological trait networks, identified either through modeling or experiment, they should *not* be viewed as “end point” ideotypes, whether they are achievable or not. Breeding programs are themselves rich sources of highly relevant trait information, much of it having been earned over many breeding cycles within and across complex target environments. Given these considerations, physiological trait networks are best used as selection criteria to *enrich* breeding programs, and only after carefully evaluating what is already known about beneficial traits, the available agronomic practices, as well as the express aims of the breeder. Integration of crop growth models with whole-genome prediction (CGM-WGP methodology) was designed to achieve this aim and is widely considered a revolution in molecular breeding (Technow et al. 2015; Messina et al. 2020). The continued development of models that enable linkage between performance, physiology, and functional genomics remain a priority for agriculture and will require the continued close collaboration of breeders, geneticists, physiologists, and modelers (Tardieu et al. 2018).

### Conclusions

We uncovered two contrasting trait networks likely to confer improved performance when water limits plant growth (particularly late-season growth) versus when water is non-limiting. These two trait networks can be understood by their aggregate effect on water use and water conservation. Dry climates with late-season deep soil water availability featured plants with conservative stomata, deep roots, and high Vpmax, whereas wet environments featured plants with risky stomata, deep roots, efficient and safe water transport, and high maximum LAI. The efficacy of these trait networks arose from climate differences among sites (precipitation amount, precipitation timing, VPD, and temperature), i.e., “envirotype” (Xu 2016). In addition to the trait differences separating these two broad water use strategies, we also found striking trait similarities *within* each of these groups (e.g., among the two “wet” and irrigated scenarios). Such generalization is important because if the benefit of a single trait network cannot be extended across multiple sites then every site and crop combination will represent an independent breeding challenge (Tardieu 2012). Custom designing crop plants for every situation is at odds with the global challenges facing agriculture. The process-based approach to crop modeling presented here may help to meet these challenges by complementing and extending site-specific experimental results to a broader range of cropping systems, soils, and climates, and thus improve our general understanding of trait network effects on water use, plant growth, and grain yield.

## Acknowledgments

The contributions of Mark Cooper, Graeme Hammer, Timothy Brodribb and Ian Wright were supported by the Australian Research Council Centre of Excellence for Plant Success in Nature and Agriculture (CE200100015). Hervé Cochard was supported by the ANR projects 16-IDEX-0001 and 18-CE20-0005. Jared Stewart was supported by the National Science Foundation (IOS-1907338).

## Data availability statement

All data and code (R, C++) used in this study are in the public domain and can be downloaded from GitHub: https://github.com/sean-gl/trait_network_ms_TREES_data_and_code

