## Supplemental Figures for "Physiological trait networks enhance understanding of crop growth and water use in contrasting environments"

**Fig. S1 Xylem vulnerability curves for embolism "susceptible" and embolism "resistant" simulations (plants). These curves represent the loss of xylem conductivity per unit decrease in xylem water potential. Thus, as the internal pressure within the xylem decreases (becomes more negative) cavitation increases and more of the conducting conduits become filled with gas, rather than water. This process starts out slowly, speeds up (steepest part of the curve) and then slows again at very low water potentials. "P50" and "P88" denote the xylem water potentials at which 50% and 88% of the maximal conductivity is lost. These points on the curves are denoted with broken (resistant) and solid (susceptible) lines and open blue symbols.**

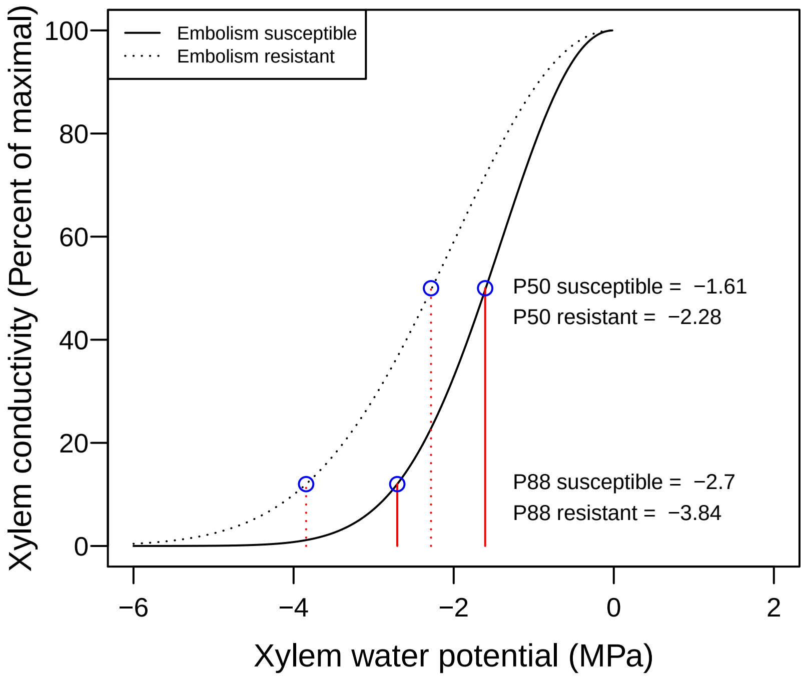

**Fig. S2** **Sap flow measured on field plants in Greeley, Colorado (orange symbols) vs simulated whole-plant transpiration (solid blue line) under fully watered conditions (100% of reference ET; left panel) and limited water (40% of maximal ET; right panel). Under full water, the $R^2$, residual standard error (RSE), and bias were 0.58, 0.655, and -0.381, whereas under limited water, the values of these fit statistics were 0.63, 0.522, and 0.111. Thus, TREES resulted in slightly more error (standardized) and bias (negative bias; underestimated transpiration) under fully irrigated conditions than under limited water. Modeled transpiration vs measured sapflow are represented as standard major axes models (inset figures).**

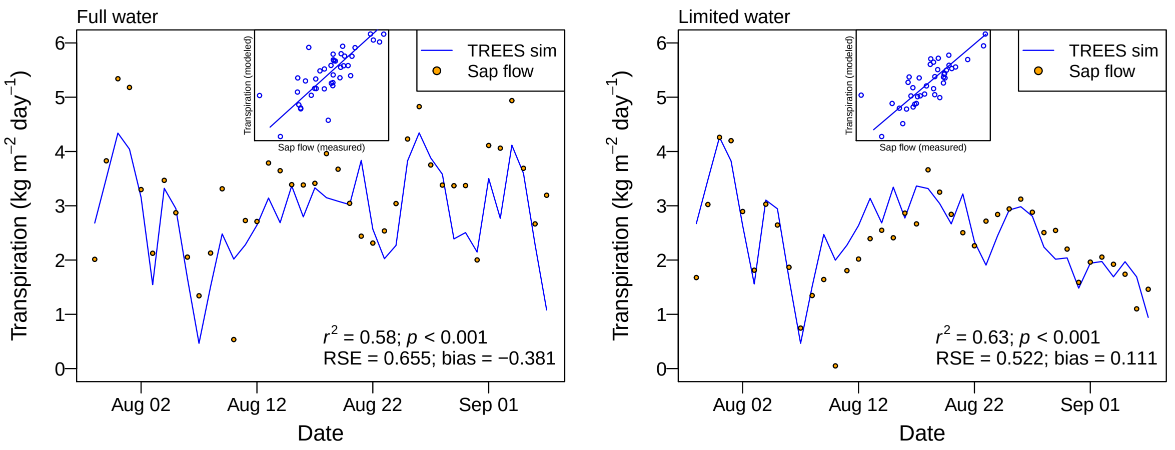

#### Net primary productivity (NPP) and reproductive output (yield) for each trait contrast (e.g., deep vs shallow rooting) for all climate scenarios. Symbol size has been scaled proportionately with variance in NPP across simulations.

##### Fig. S3 Central Plains, Wet

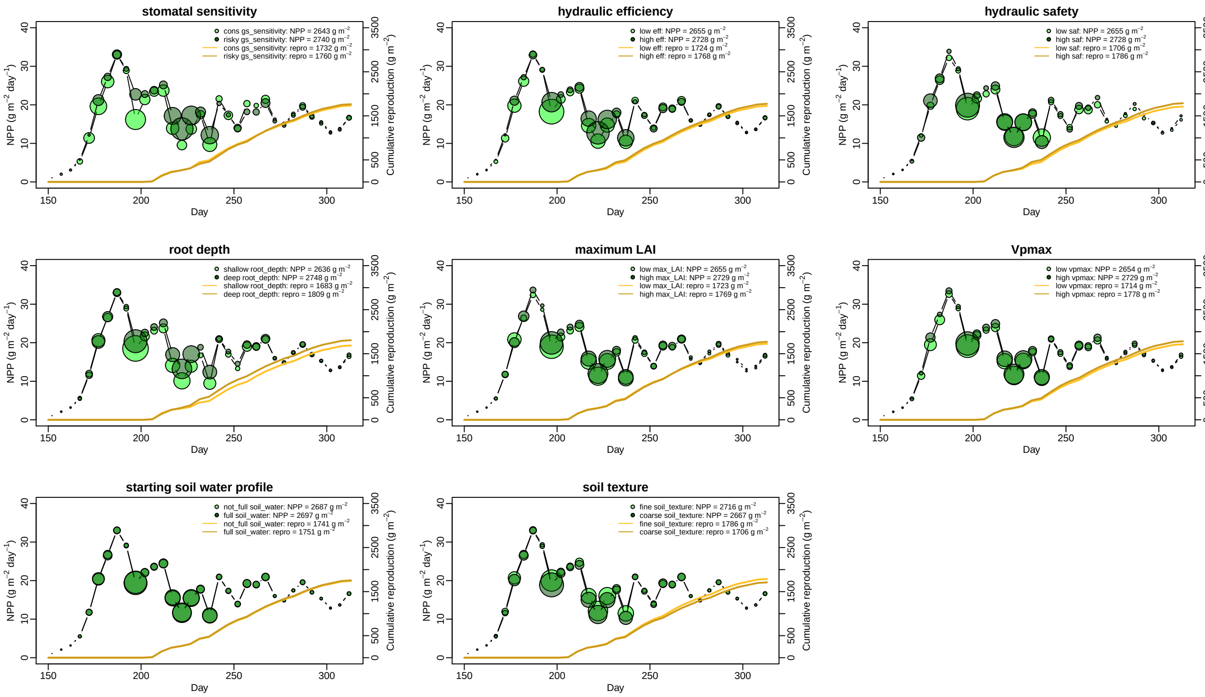

### Fig. S4 Central Plains, Dry

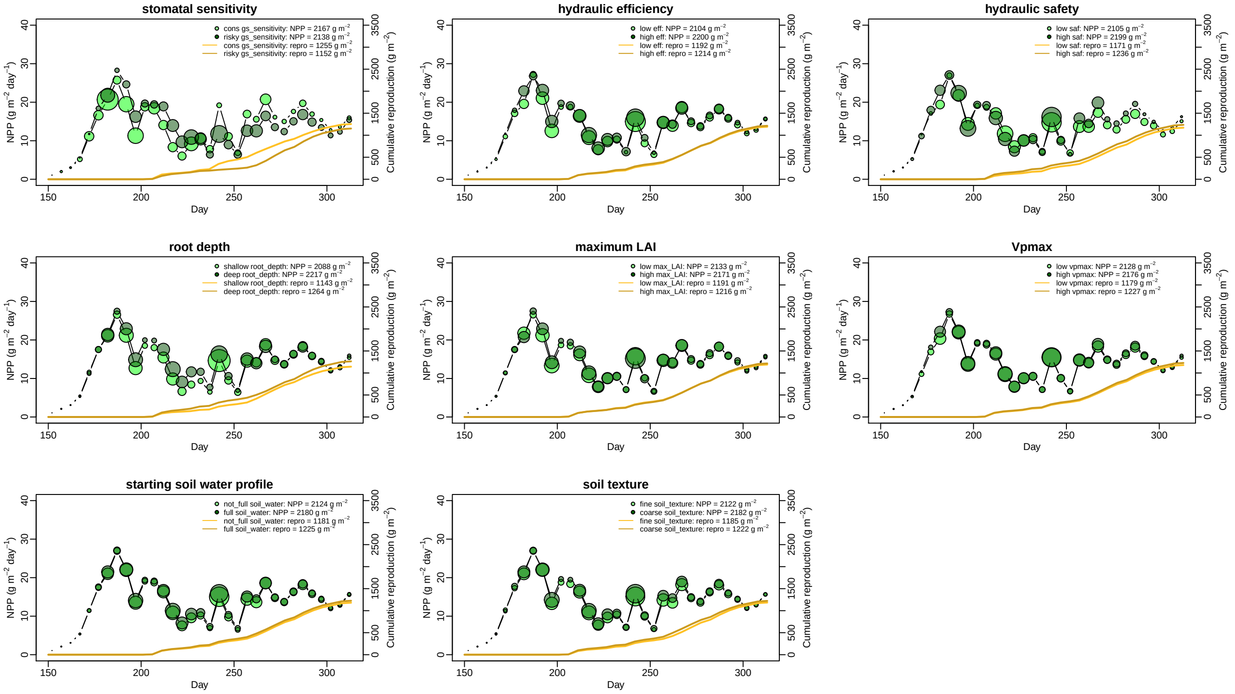

### Fig. S5 High Plains, Wet

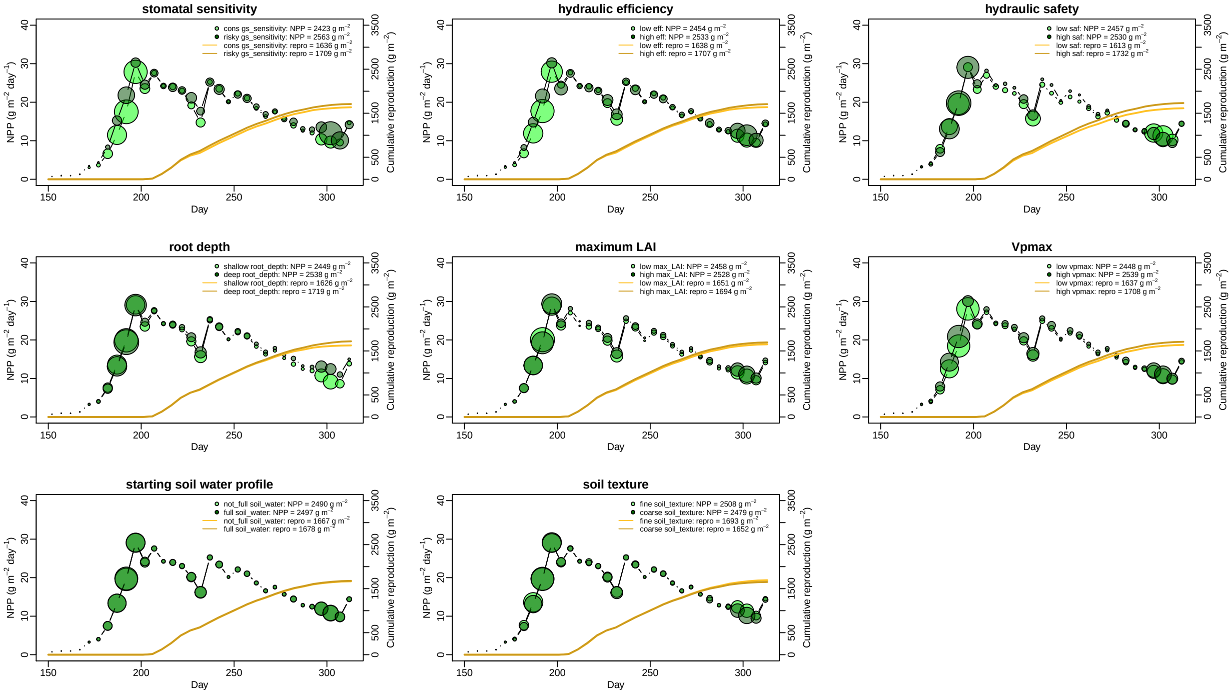

### Fig. S6 High Plains, Dry

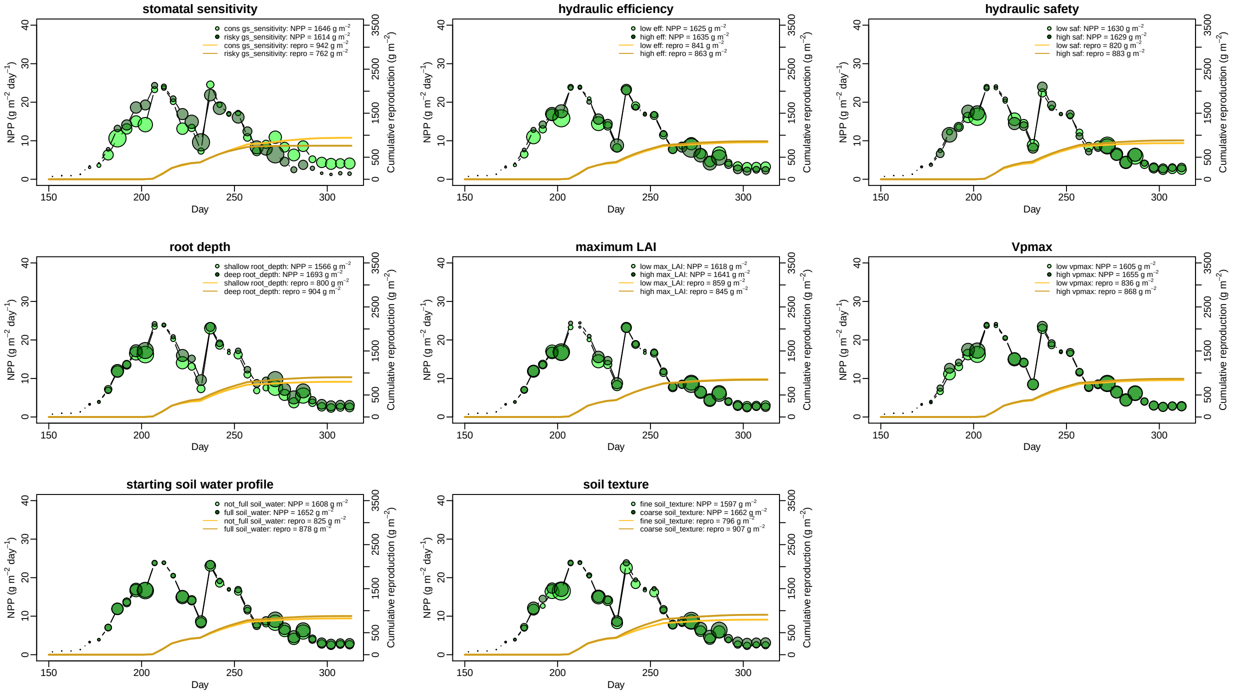

### Fig. S7 Irrigated

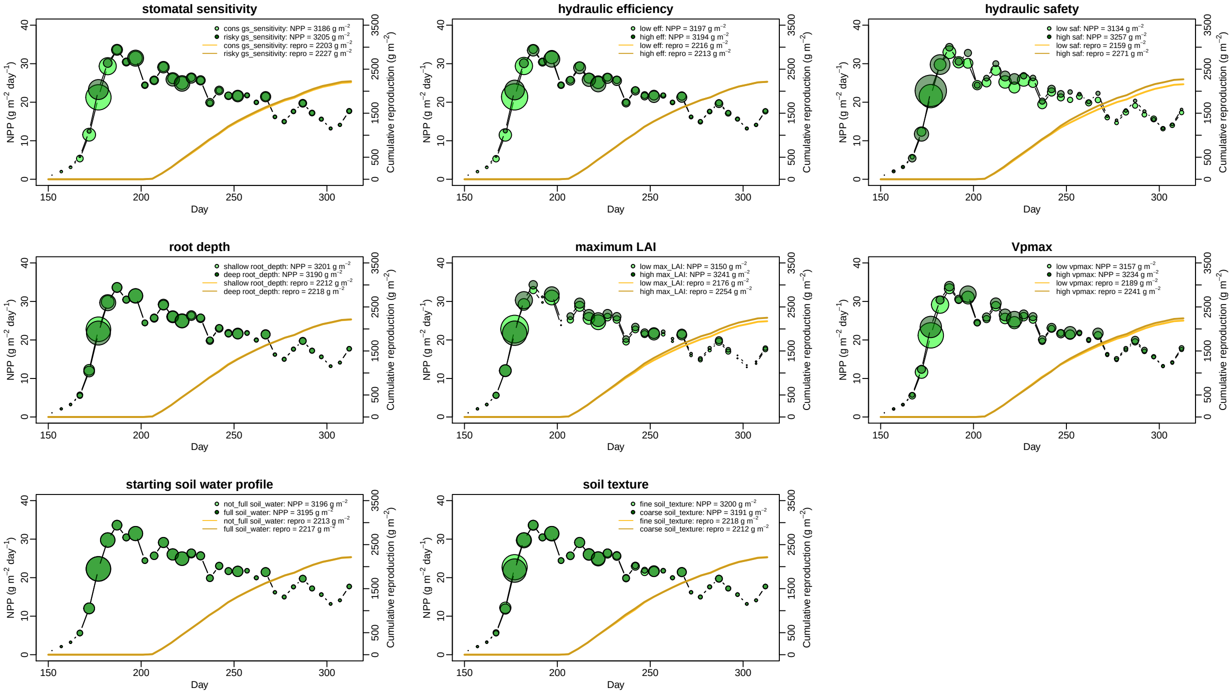

**Root area and soil water content for shallow (0-5 cm), intermediate (15-35 cm), and deep (75-105 cm) soil layers for each treatment and each trait contrast (e.g., "risky" vs "conservative" stomata). Each page represents a different climate scenario.**

**Fig. S8 Central Plains, Wet**

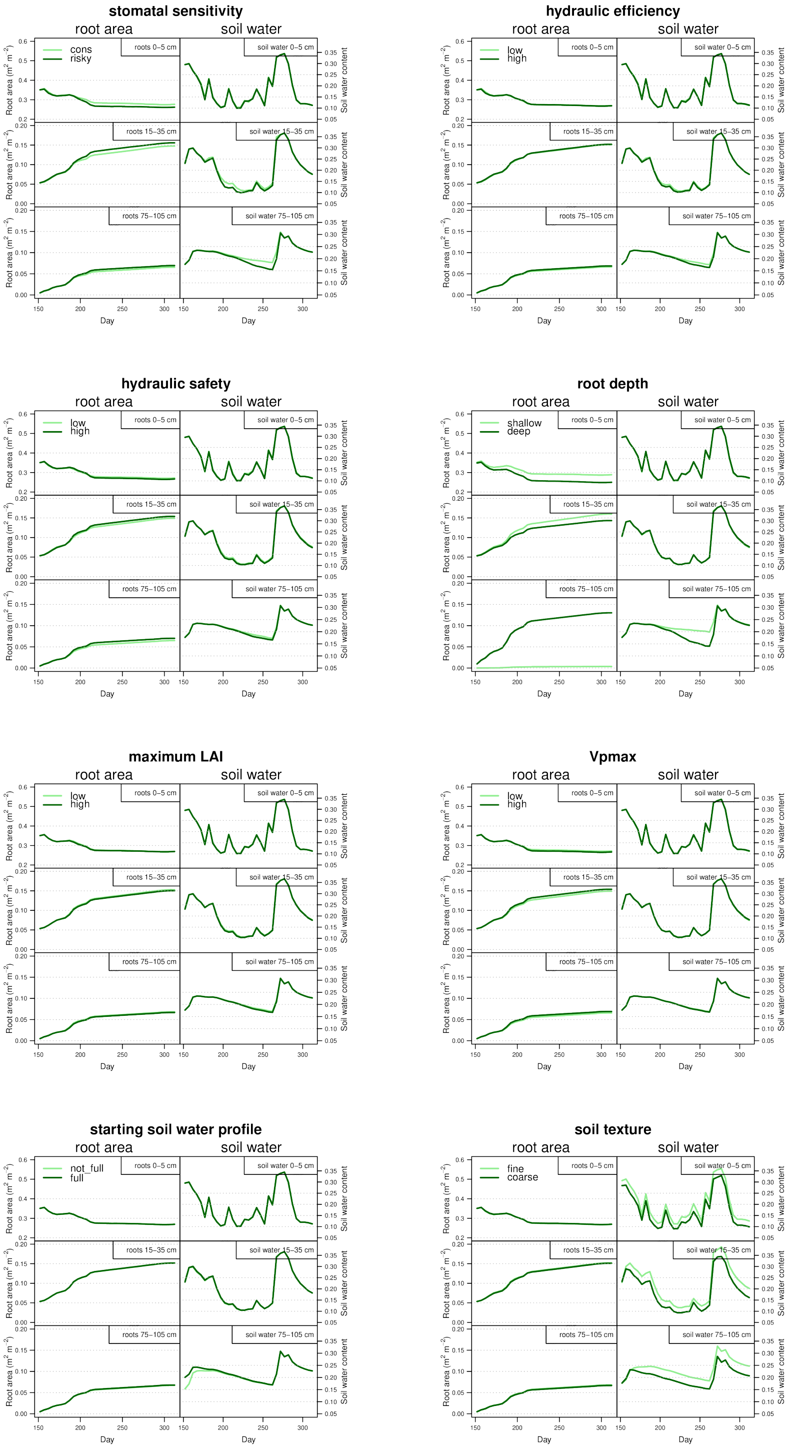

### Fig. S9 Central Plains, Dry

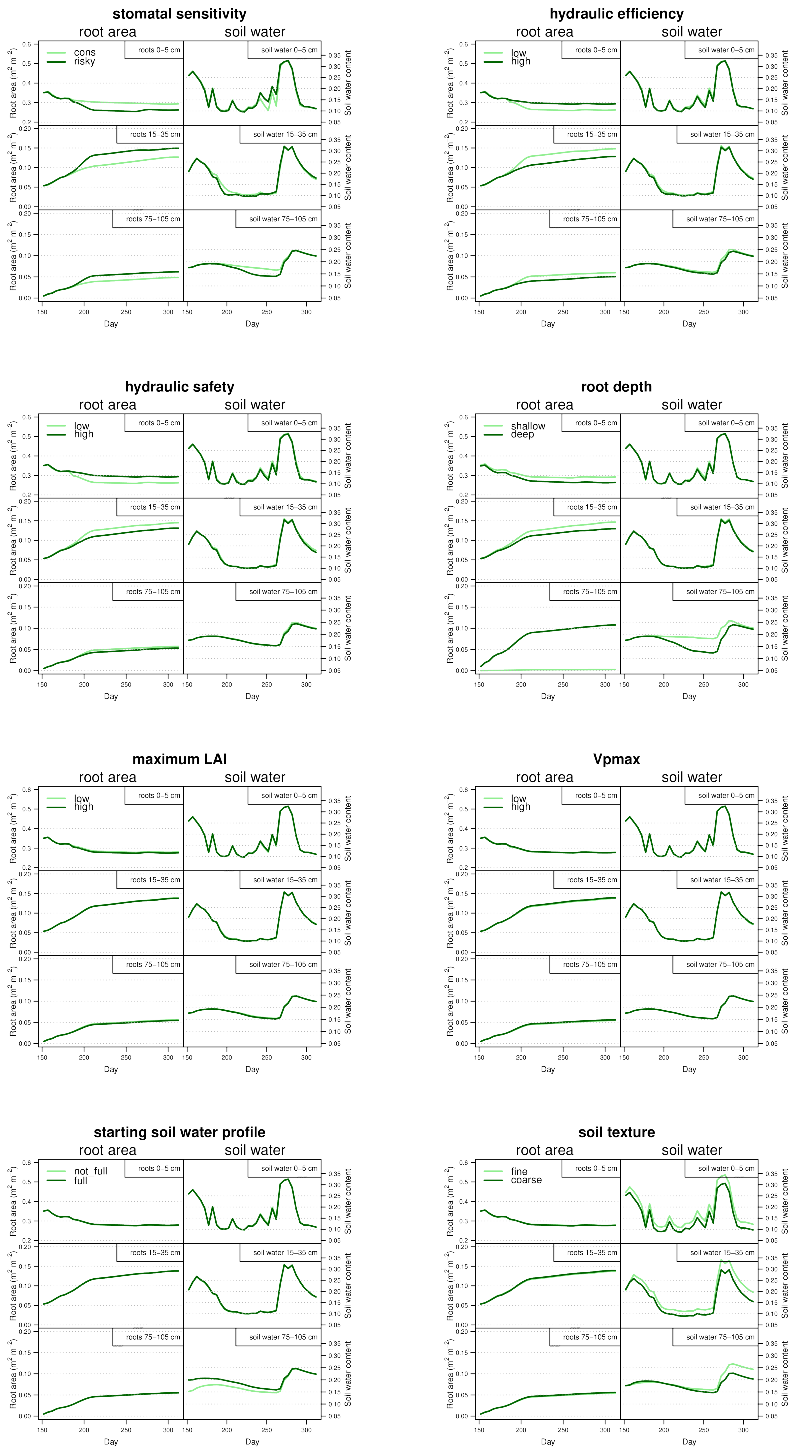

### Fig. S10 High Plains, Wet

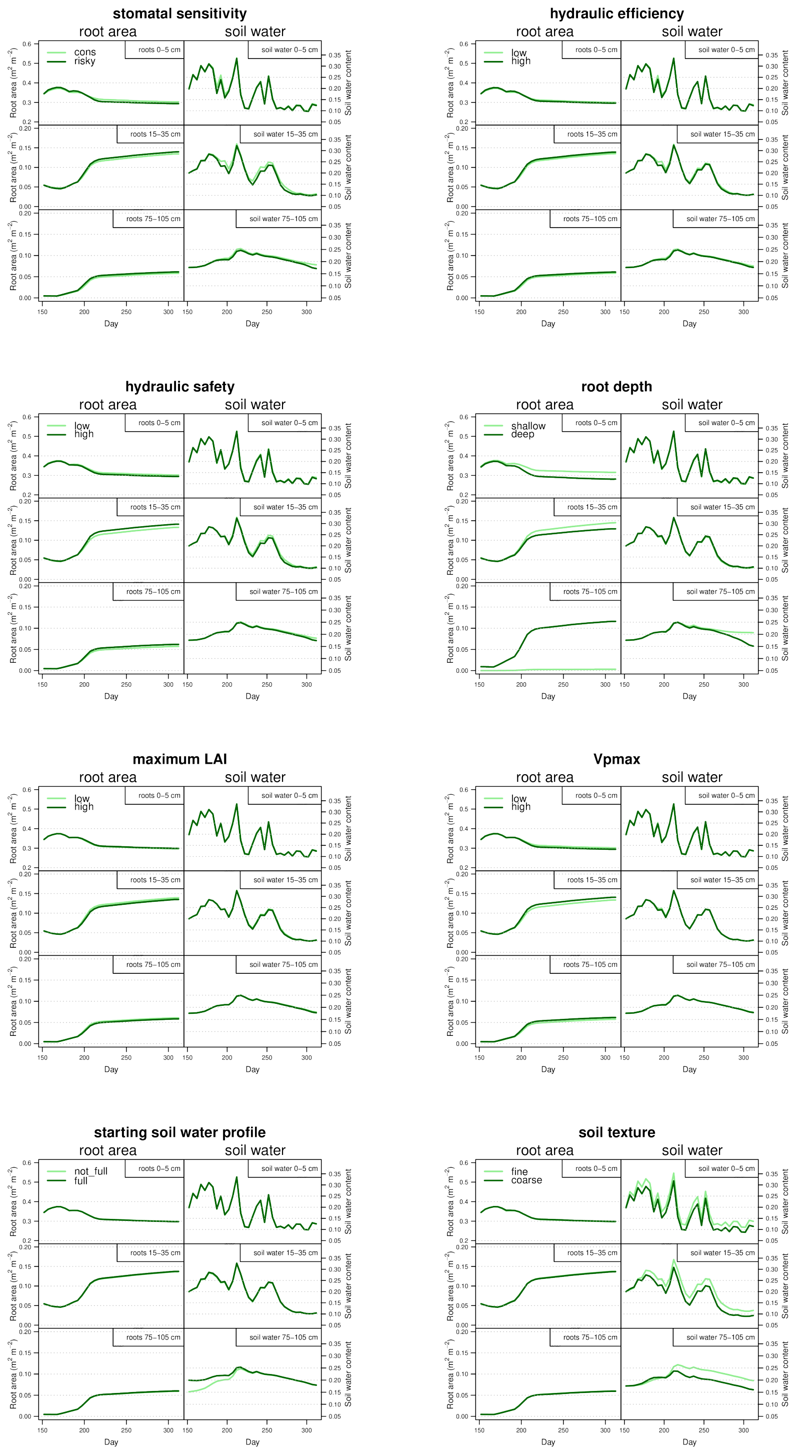

### Fig. S11 High Plains, Dry

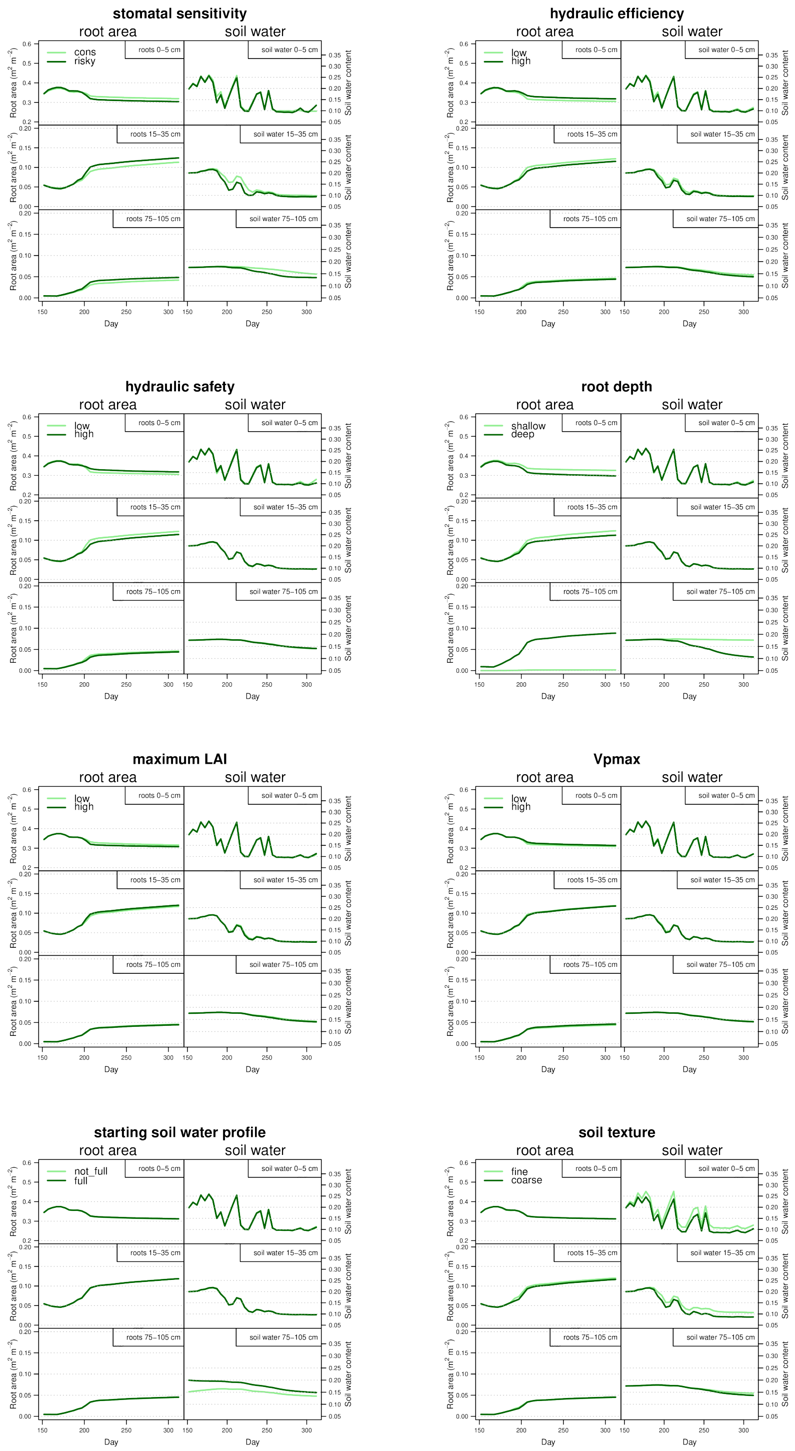

#### Fig. S12 Irrigated

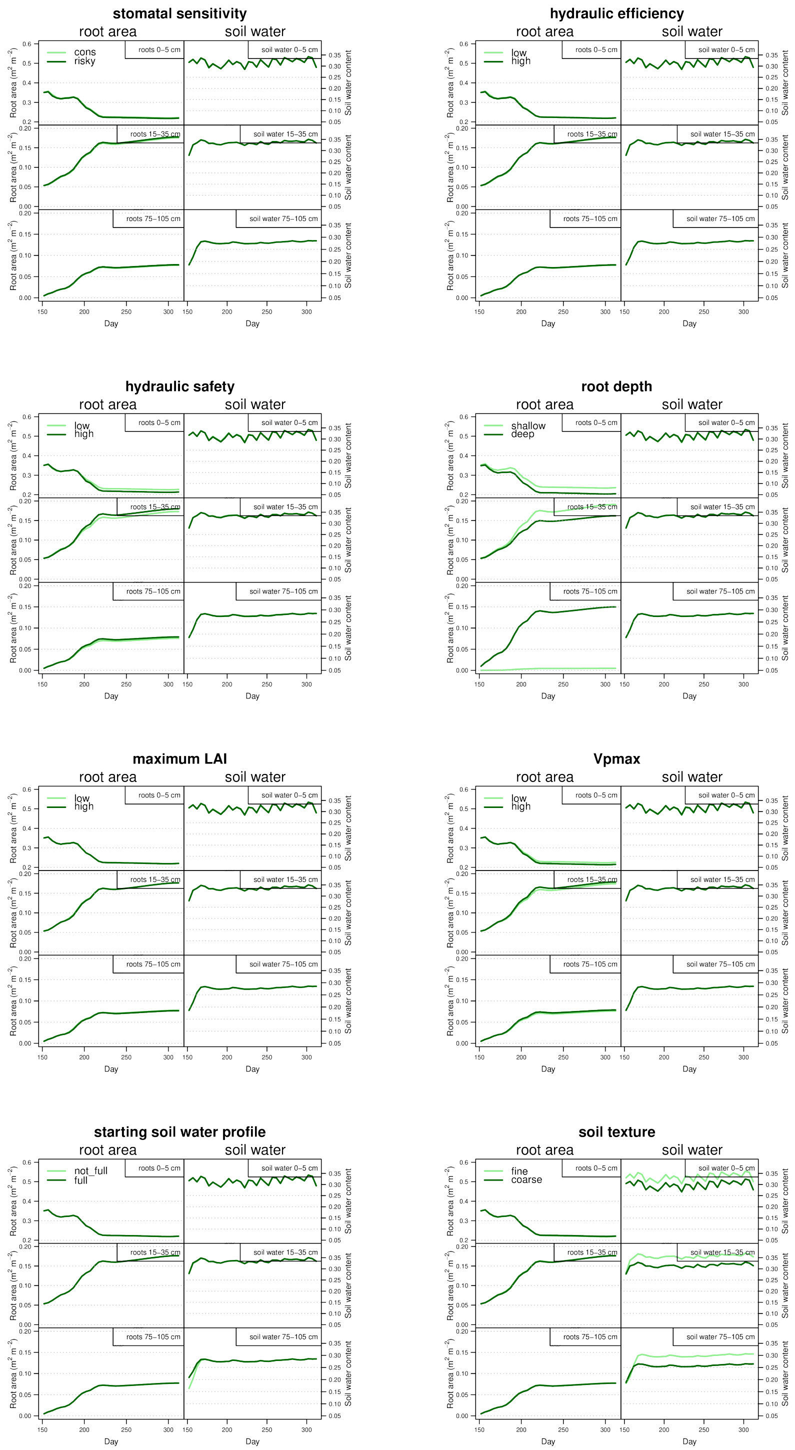

**Xylem and soil conductances (stacked bars) and whole-plant transpiration for each trait contrast. Each page represents a different climate scenario. "T_fraction" = the fraction of total seasonal precipitation that was transpired. "PrUE" = total seasonal NPP / total seasoanl precipitation.**

**Fig. S13 Central Plains, Wet**

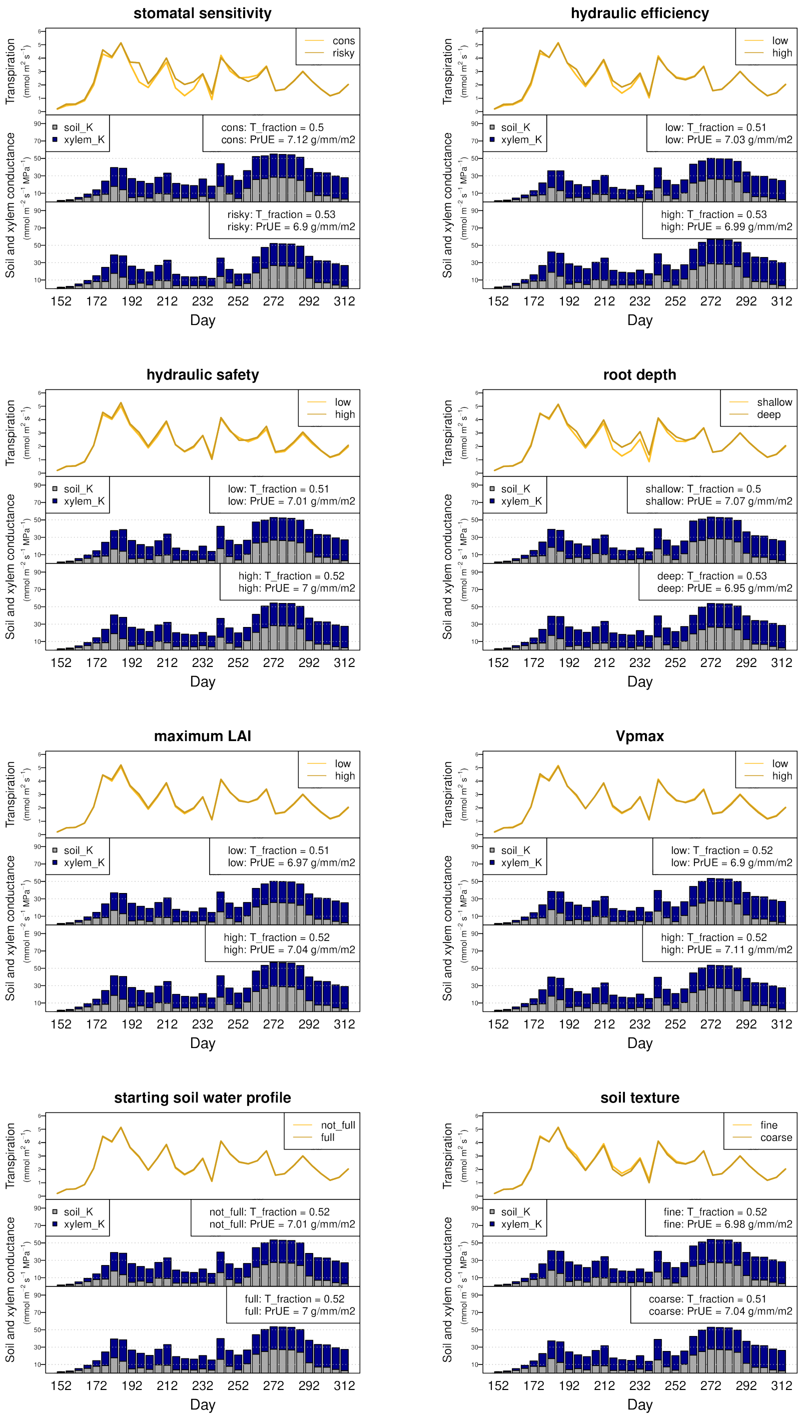

### Fig. S14 Central Plains, Dry

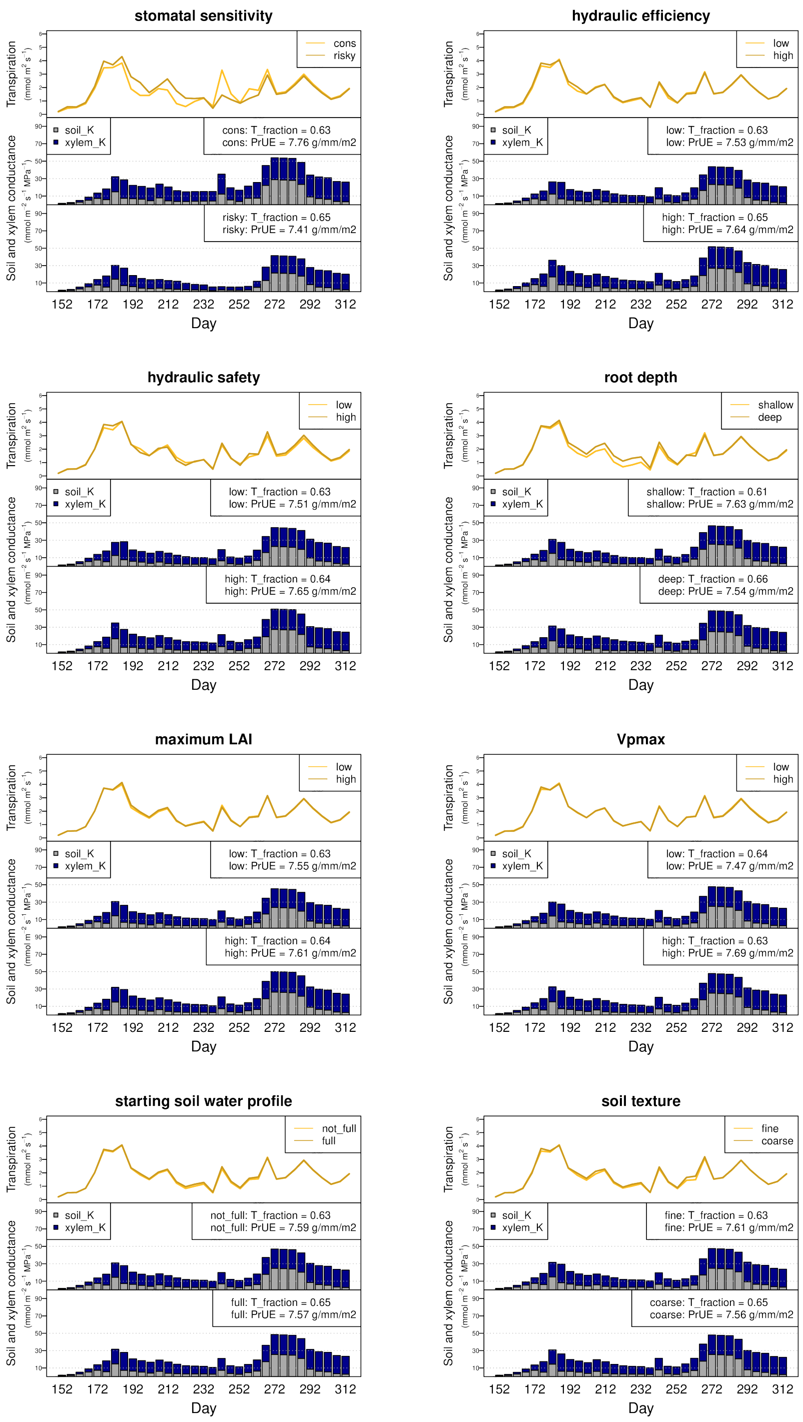

### Fig. S15 High Plains, Wet

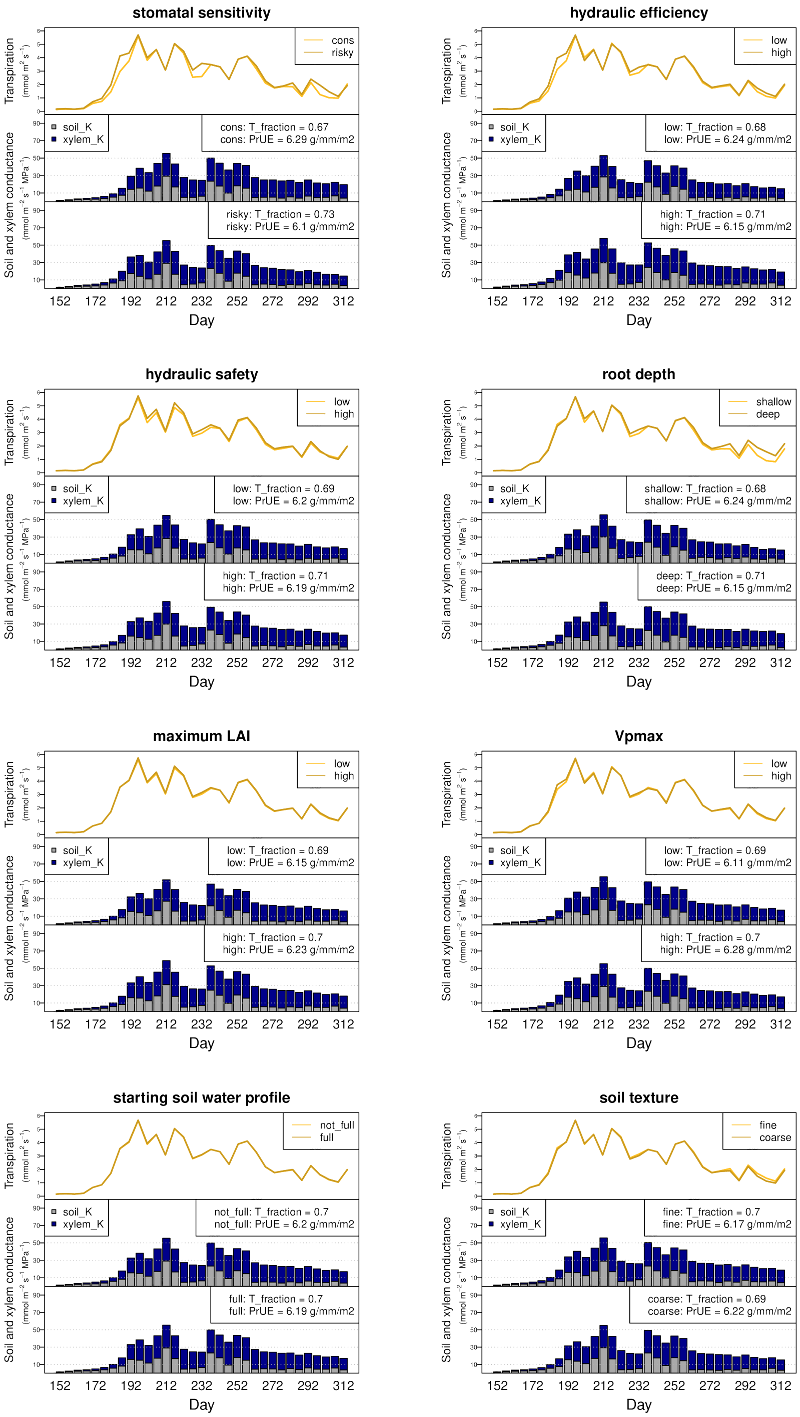

### Fig. S16 High Plains, Dry

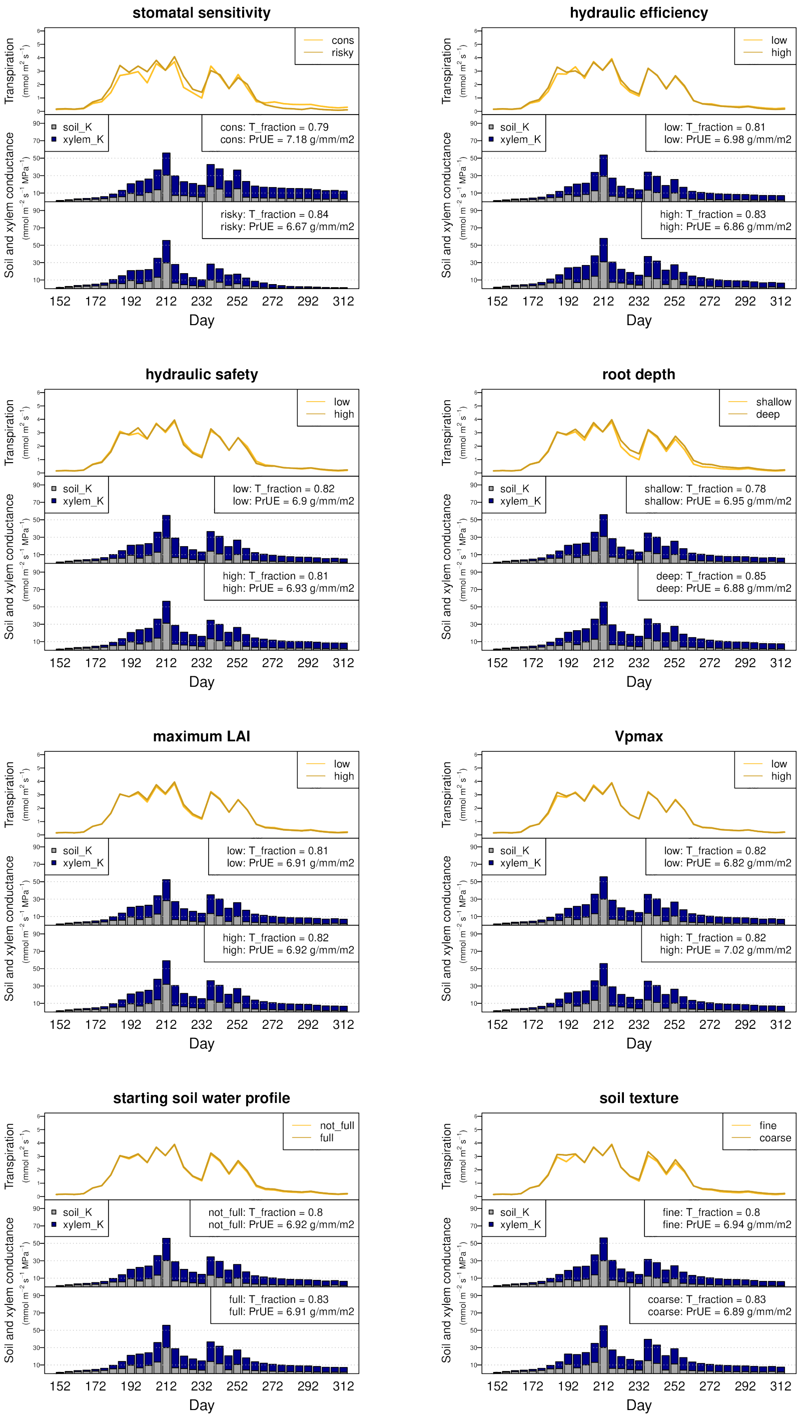

### Fig. S17 Irrigated

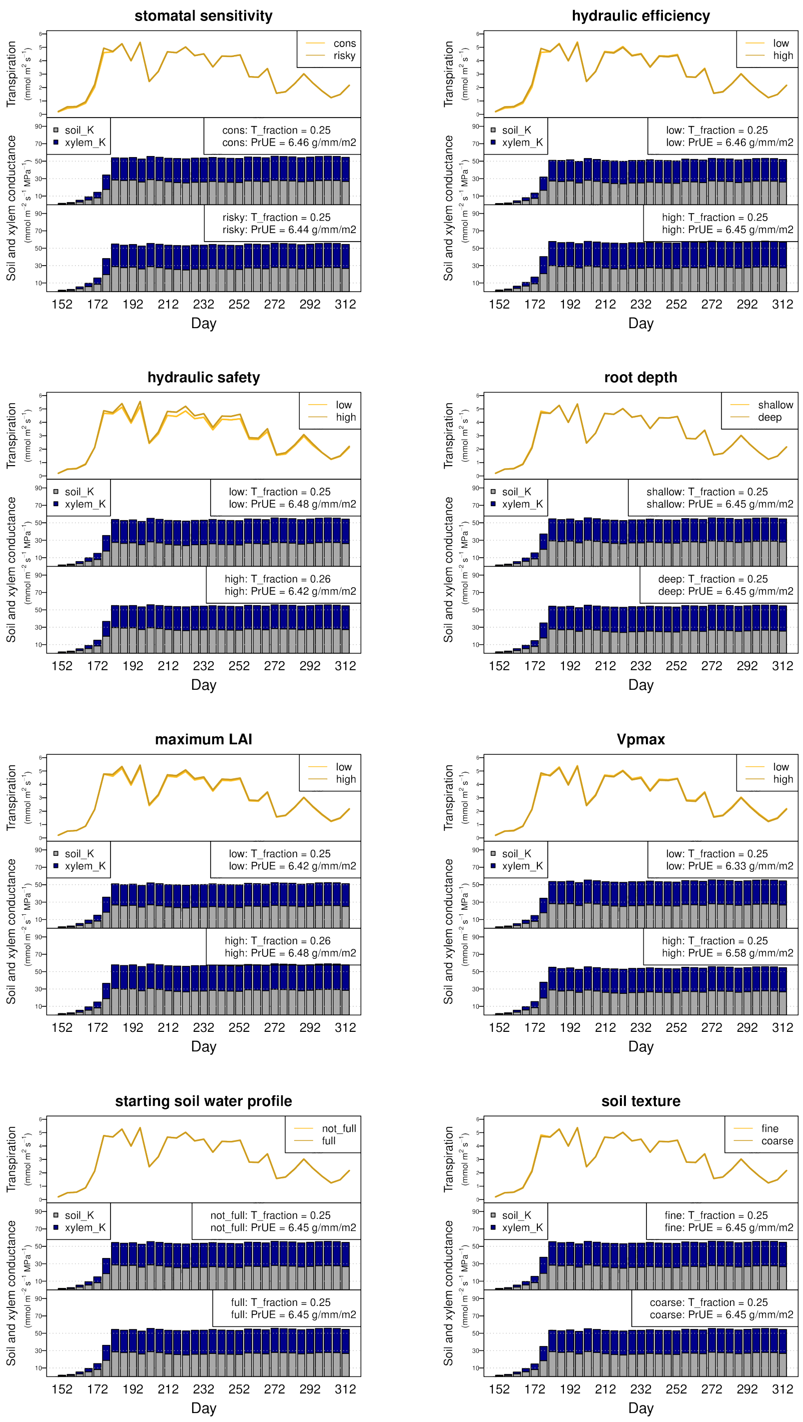

**Representative multiple trait decision trees for all climate scenarios. The first branch point (trait contrast) is the trait resulting in the largest decrease in model error, whereas the last branch point denotes the trait contrast resulting in the smallest decrease in model error. Decision trees were drawn only four nodes, thus representing the four most "important" traits. Error bars denote +/- 1 standard deviation (n=16). Vpmax = maximum activity of PEP-carboxylase. Root depth = maximum depth of root system. Max LAI = maximum achievable leaf area index. Saf = xylem embolism resistance. Gs sensitivity = stomatal response to leaf water potential. Eff = maximum xylem conductance.**

**Fig. S18 Central Plains, Wet**

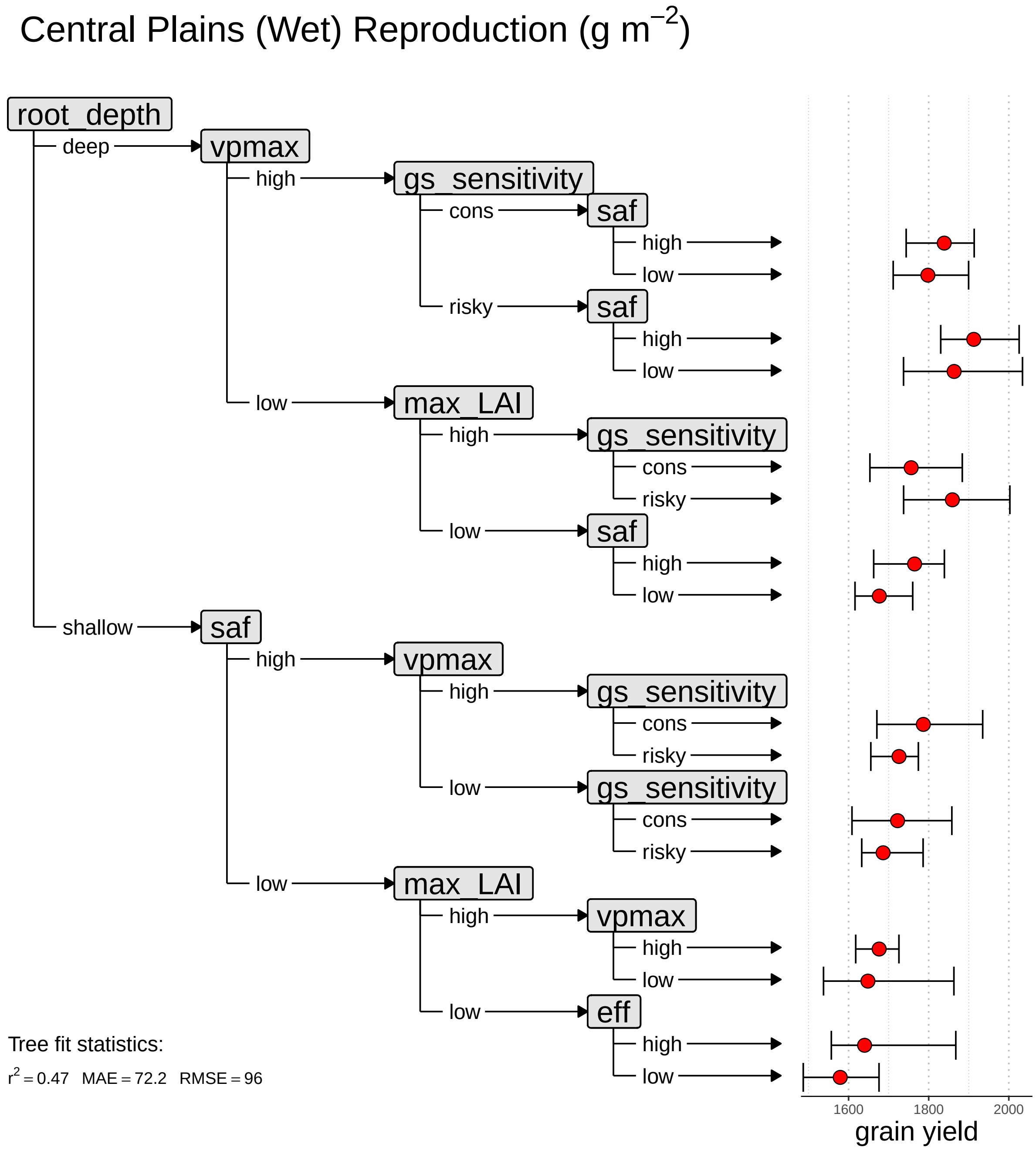

**Fig. S19 Central Plains, Dry**

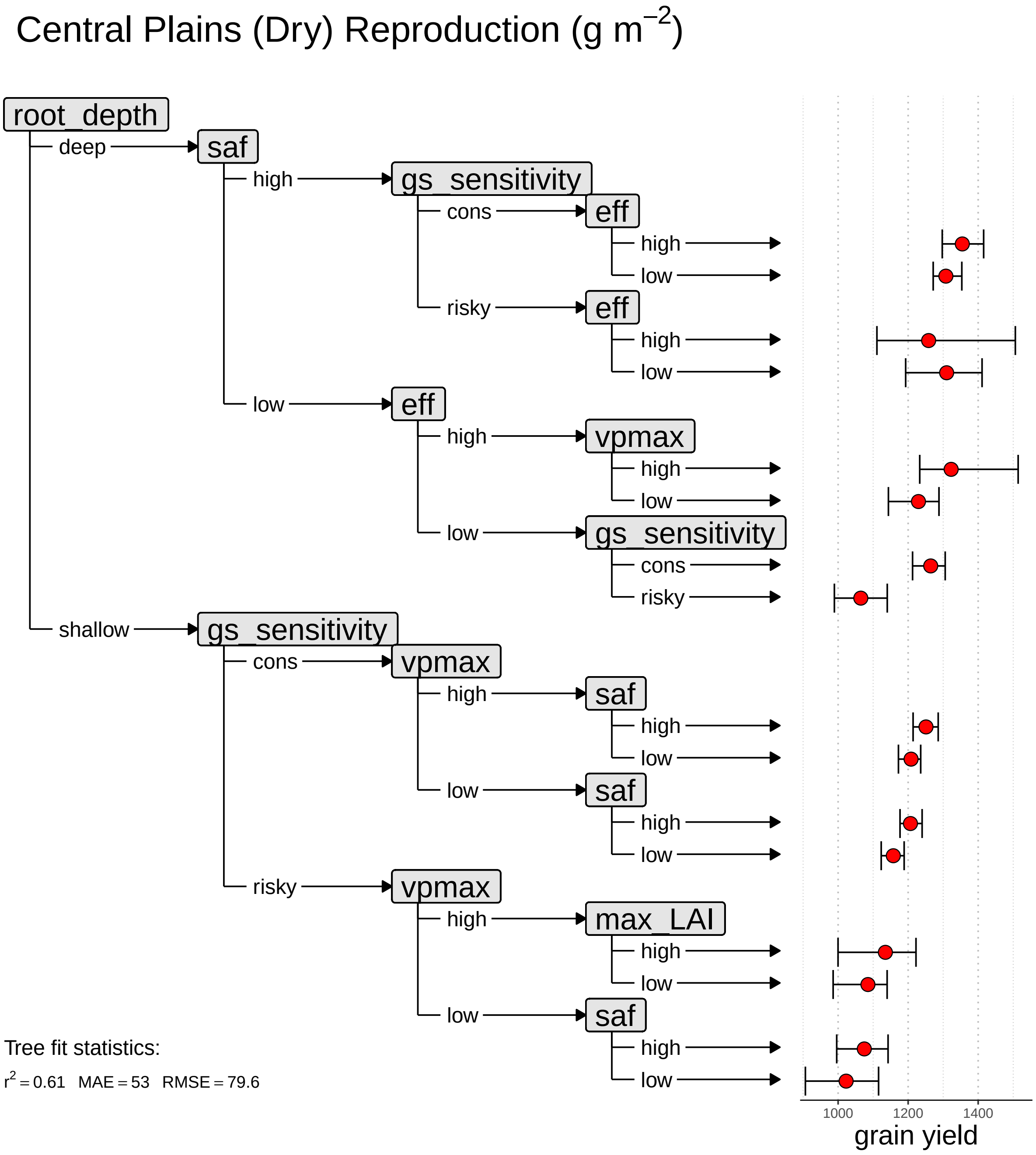

**Fig. S20 High Plains, Wet**

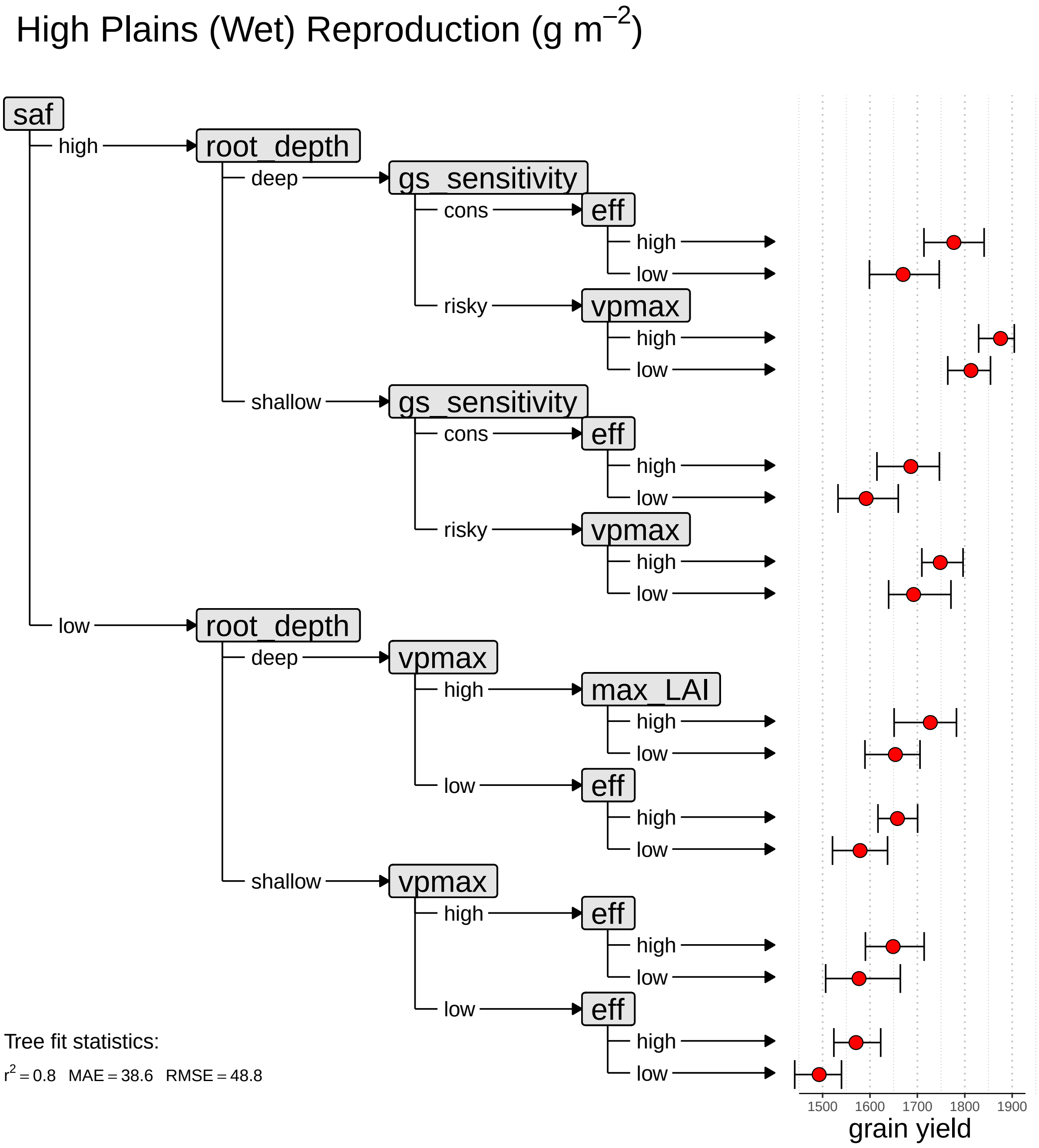

#### Fig. S21 High Plains, Dry

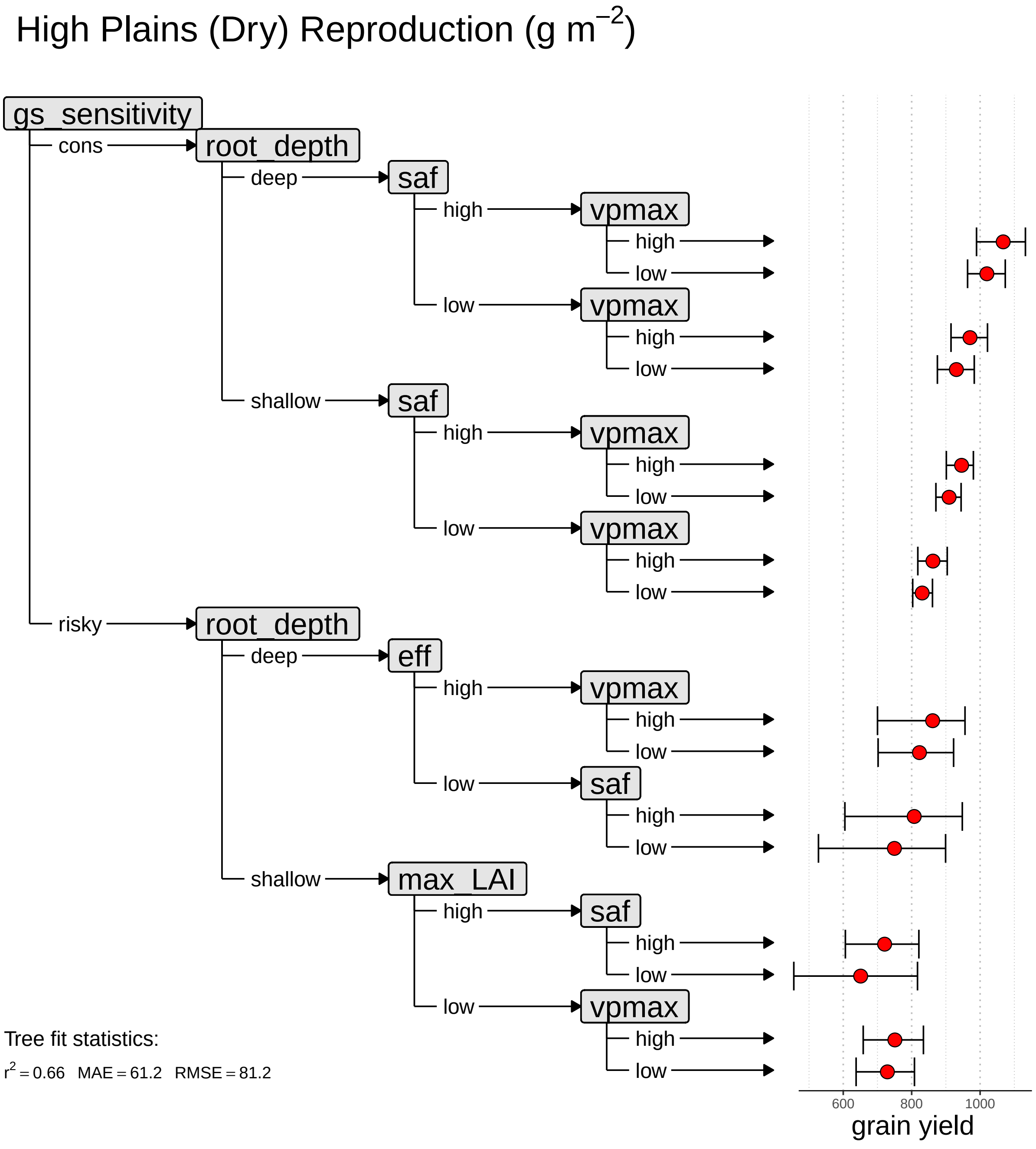

**Fig. S22 Irrigated**

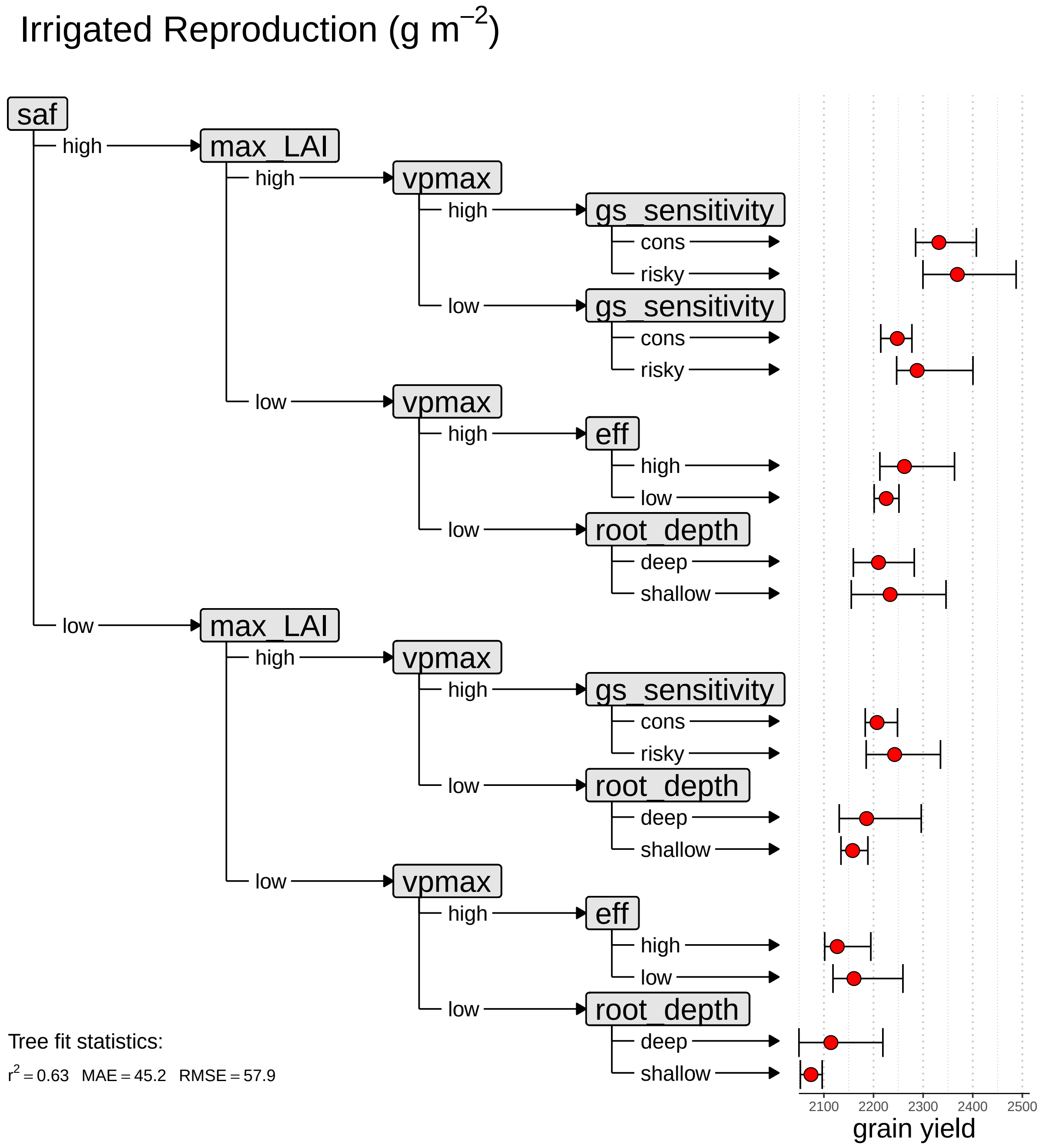

**Reproductive output heatmap for all climate scenarios. Each square represents the reproductive output of two "stacked" plant and/or soil traits. Color intensity denotes reproductive output. Vpmax = maximum activity of PEP-carboxylase. Root depth = maximum depth of root system. Max LAI = maximum achievable leaf area index. Saf = xylem embolism resistance. Gs sensitivity = stomatal response to leaf water potential. Eff = maximum xylem conductance. Soil water = presence or absense of soil water in deepest layer. Soil texture = texture of entire soil column.**

**Fig. S23 Central Plains, Wet**

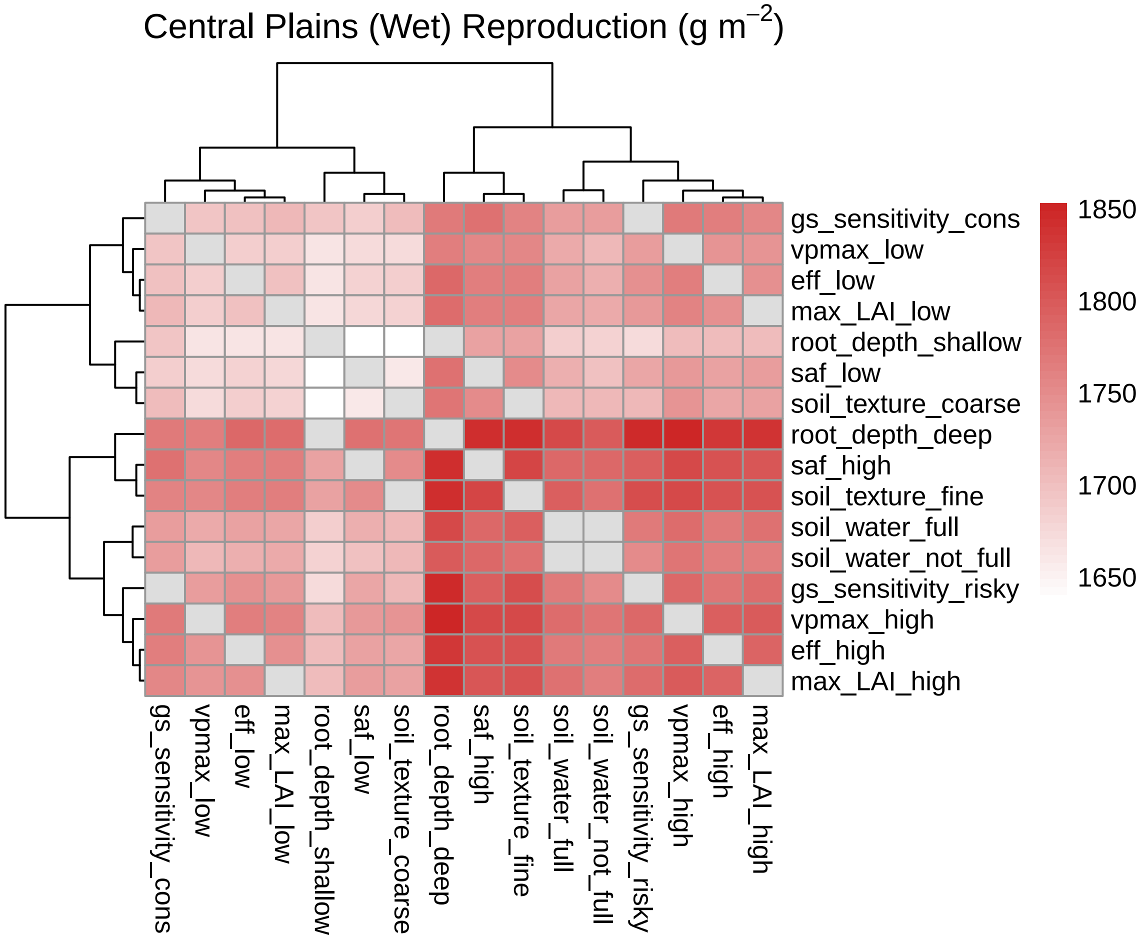

**Fig. S24 Central Plains, Dry**

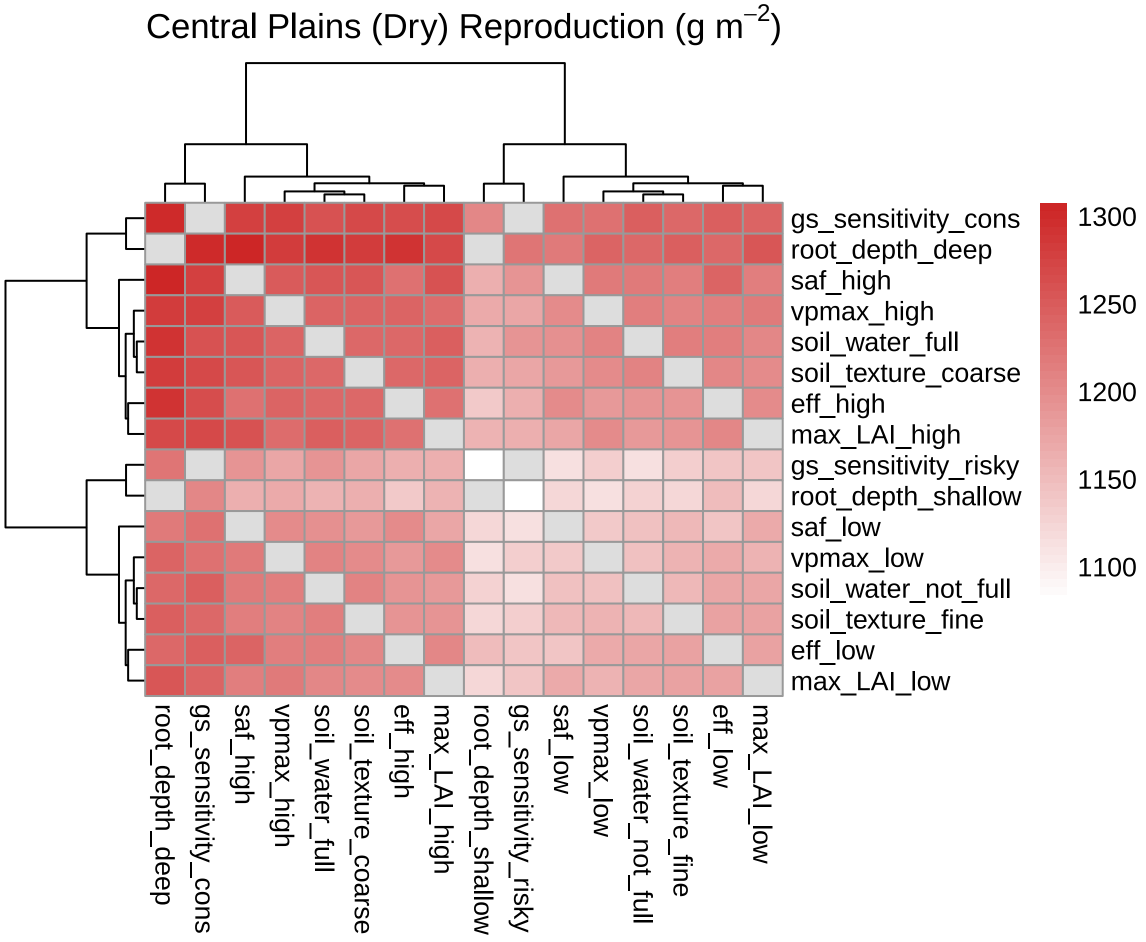

**Fig. S25 High Plains, Wet**

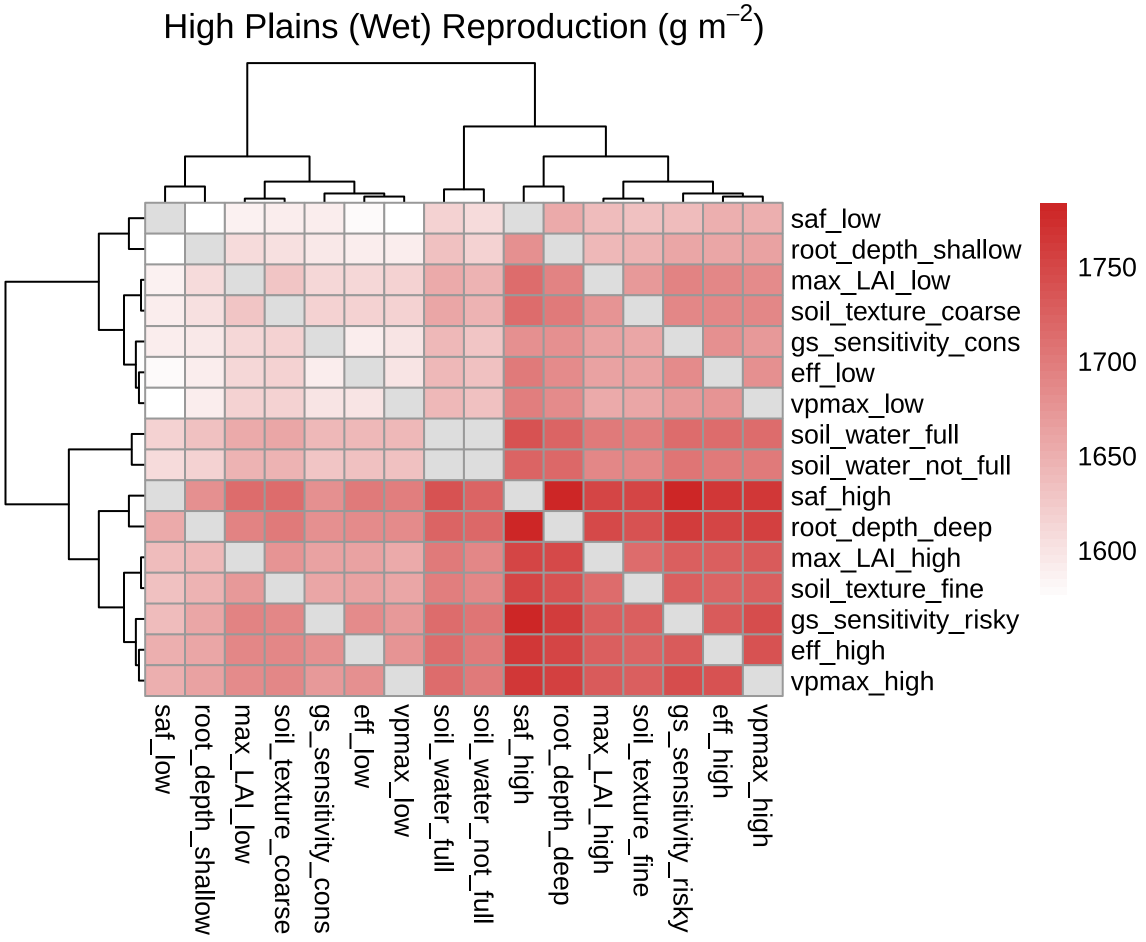

**Fig. S26 High Plains, Dry**

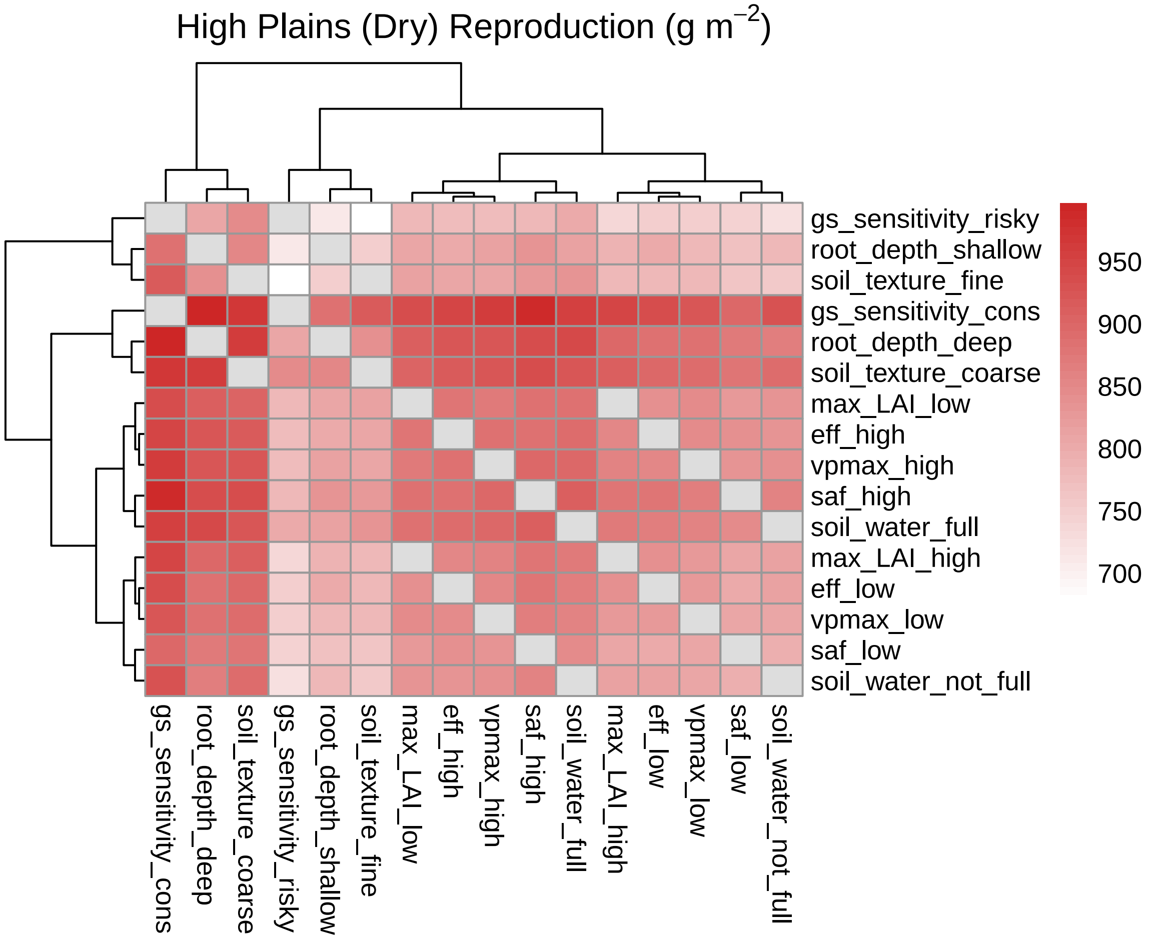

**Fig. S27 Irrigated**

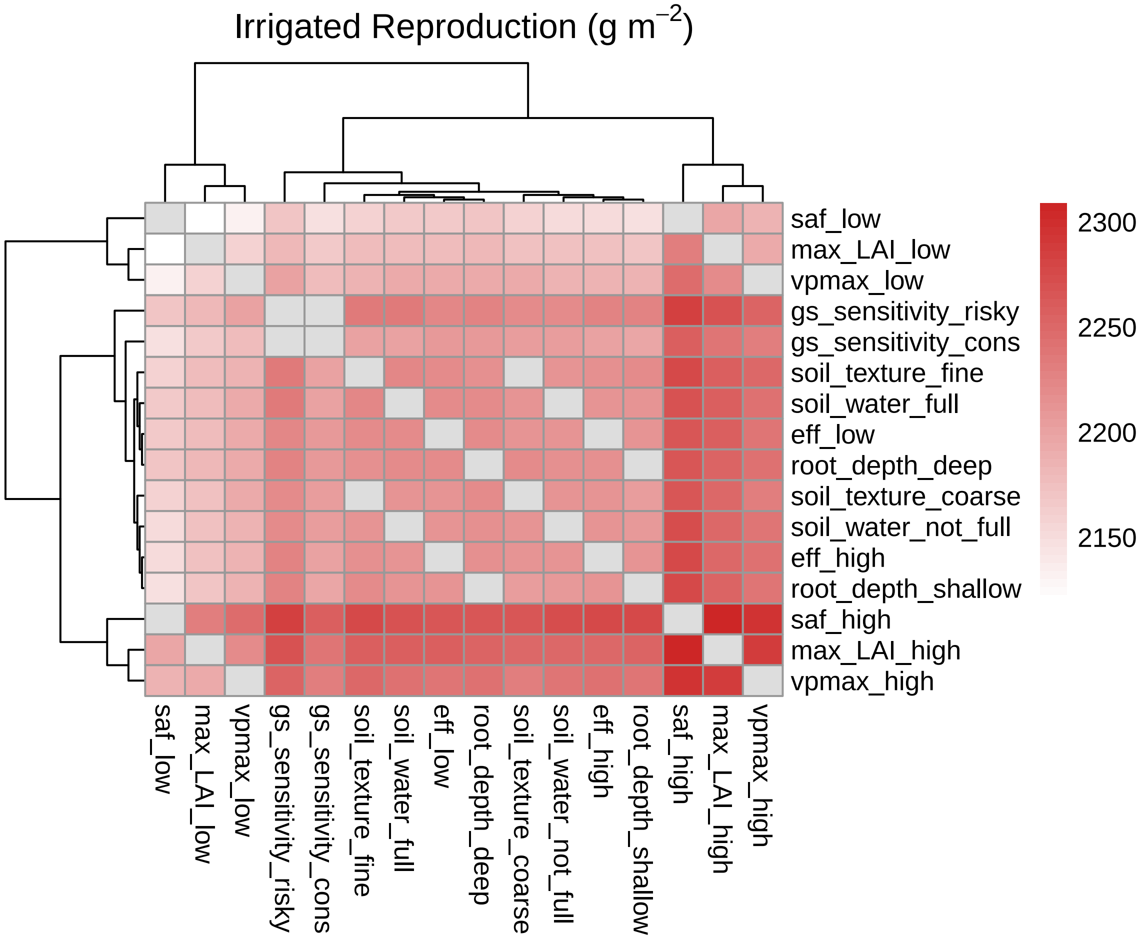
